# flexLiTE: flexible micro-LED integrated optoelectrodes for long-term chronic deep-brain studies

**DOI:** 10.1101/2022.08.05.503006

**Authors:** Eunah Ko, Jose Roberto Lopez Ruiz, Mihály Vöröslakos, Meng-Lin Hsieh, György Buzsáki, Euisik Yoon

**Affiliations:** Department of Electrical Engineering and Computer Science, University of Michigan, Ann Arbor, MI 48109, USA; Neuroscience Institute, Langone Medical Center, New York University, New York, NY 10016, USA; Department of Biomedical Engineering, University of Michigan, Ann Arbor, MI 48109, USA; Department of Mechanical Engineering, University of Michigan, Ann Arbor, MI 48109, USA; Center for Nanomedicine, Institute for Basic Science (IBS) and Graduate Program of Nano Biomedical Engineering (Nano BME), Advanced Science Institute, Yonsei University, Seoul 03722, South Korea

## Abstract

Understanding complex neuronal circuitry and its functions of a living organism requires a specialized tool which is capable of recording a large ensemble of neuronal signals at single cell resolution and modulating neuronal activities selectively in the target region of brains with high spatiotemporal resolution, while sustaining long-term chronic operation without significant tissue degeneration or device shifts. We hereby present an ultra-flexible, minimally-invasive, Michigan-type neural probe for chronic opto-electrophysiology studies: flexLiTE (flexible micro-LED integrated optoelectrodes). flexLiTE incorporates monolithically integrated, soma-sized inorganic micro-LEDs (12 individually operated) and 32 recording electrodes. Both stimulation and recording modalities were achieved by stacking two modules on a flexible substrate: one with micro-LEDs for neuromodulation and the other with recording sites, resulting in a 115 μm-wide,12 μm-thick, 10 mm-long optoelectrode. From prototype devices, we demonstrated the reliable operation of flexLiTEs for recording and modulation of hippocampal neurons in a freely moving mice for over ∼8 month.

## Introduction

Recent advances in micro-fabricated deep-brain implantable neural interfaces unraveled circuit connectivity of neurons to elucidate functions of individual neurons as well as give a hint of mechanisms behind neurological disorders for better treatment [1–4]. A large number of neurons could be simultaneously recorded while precision neuromodulation was enabled by delivering light onto target neurons for optogenetic stimulation [5–7]. Yet, long-term (weeks to months) recording along with optogenetic stimulation remains as a key challenge in implantable probes for deep-brain regions. The penetration volume and the mechanical stiffness should be reduced to prevent insertion damage, foreign-body induced tissue reaction and following fibrotic scarring from micro motions, which degrades quality of signals and control of cells over time [8–9].

Flexible substrates can alleviate the mechanical stiffness and tissue damage compared to rigid silicon probes [10–11]. The stiffness of a probe shank can be defined as *Ewt^3^/4L^3^*, where *E* is the Young’s modulus of the shank material, *w* is the width of the shank, *t* is the thickness, and *L* is the total length. Polymer-based flexible substrate can give a low Young’s modulus of <10 GPa and can be easily realized in a relatively thin layer of < a few microns. Therefore, flexible polymer-based neural probes are beneficial for less tissue reaction and minimally-invasiveness for long-term chronic recording [12–13].

Among many flexible polymer materials, such as parylene-C, SU-8, and polyimide [14–17], recent studies showed that polyimide (PI) has excellent microfabrication and bio compatibility [18–23]. It can withstand thermal processes lower than its glass transition temperature of 360 °C [24], and can be easily micro-patterned using dry reactive ion etching (RIE). Recent progress demonstrated that the high-density recording electrodes could be integrated on a polyimide substrate for reliable recording of spikes in a large-population of cells [25–26].

Even though the stability and longevity of the polyimide-based flexible neural probes has been extensively studied, less progress has been done for integration of perturbation sources. Among many neuromodulation methods, optogenetics is one of the most widely used ones by virtue of its specificity and high spatiotemporal controllability. Genetically modified opsins that are present on the surface of cell membranes are activated by light at a specific wavelength, leading to neuronal spiking or inhibition [27–31]. Optogenetic light source should suit for minimally-invasive flexible neural interface and the requirements can be summarized as follows: (i) light shall illuminate and perturb only a small group of cells with a high spatiotemporal resolution; (ii) footprint should be small enough to fit the volume that is similar to the state-of-the-art flexible recording probes (∼100 µm wide, ∼10 µm thick); and (iii) each light source should be controlled individually and confined only onto the target region of the brain without illuminating light to non-targeted areas.

Micro-sized inorganic light-emitting-diodes (µILED) were first monolithically integrated onto a silicon shank [32]. Wu et al. demonstrated the first silicon optoelectrode by monolithically integrating an array of individually operated, soma-sized (10 µm x 15 µm) µILEDs on a typical Michigan-type multi-shank neural probe configuration [32]. Compared to other light delivery schemes, such as optical fibers, multi-photon stimulation, or waveguide technologies [33–37], this µILED optoelectrode has many advantages including high scalability, fabrication process compatibility (mass-production by utilizing wafer-level manufacturing), and high temporal resolution for selective neuromodulation inside the deep-brain region [38–39].

In this work we present an ultra-flexible Michigan-type optoelectrode by monolithically integrating cell-sized µILEDs into a polyimide substrate. This approach is different from previous ones [40–41] in a way that we monolithically integrate the small soma-size µILEDs (∼10 µm) through wafer-level processing instead of manual die transfer of individual LEDs which is limited to a relatively large size (> ∼50 µm). Our first prototype optoelectrode incorporated 12 µILEDs onto a 115 μm-wide, 6 μm-thick and 10 mm-long polyimide probe shank. With the µILEDs that span 1 mm at the tip side of the probe, flexLiTE can induce optogenetic stimulation while recording single neuron activities when paired with recording sites precisely located in proximity of µILEDs. For this, a separate recording module with 32 recording electrodes (13 µm x 15 µm) was fabricated on a flexible substrate [25] and assembled with the µILEDs module. An interposer was instrumental for the assembly where the stacked modules were ball bonded with precision alignment. We demonstrated the long-term chronic in-vivo optogenetic experiments from the fabricated flexLiTE in a freely moving mouse.

## Results

### Modularity of flexLiTE optoelectrodes

The flexLiTE explores the feasibility of a polyimide-based flexible optoelectrode integrated with micron-sized ILEDs for chronic optogenetic stimulation and recording (Fig. 1A). Two modules (a blue-light illuminating stimulation module and a recording module) are integrated into one device by self-aligned precision stacking, which is explained in detail in the Method section (Fig. 1B-E). The flexLiTE is highly flexible since it is made of polyimide with a total thickness of 6 µm for each module (Fig. 1C). The device is composed of several parts: (i) a stimulation module where the blue-ILEDs are monolithically integrated on a flexible polyimide substrate for optogenetic stimulation, (ii) a recording module where an array of electrodes are fabricated on a flexible polyimide substrate for recording wide-band neuronal signals, both spikes and local field potentials (LFPs), (iii) an interposer that is being used for stacking and aligning two modules ((i) and (ii)), (iv) a custom-made cable that connects the interposer to a headstage, and (v) a headstage PCB with an Omnetics connector. All the components are assembled into a complete flexible optoelectrode microsystem (Fig. 1D). Each module, stimulation or recording, is separately fabricated and tested before assembly, which will enhance the final assembly yield. The 12 µILEDs can be individually controlled, either activated at the same time or individually activated in sequence (Fig. 1B and E).

**Fig. 1.**
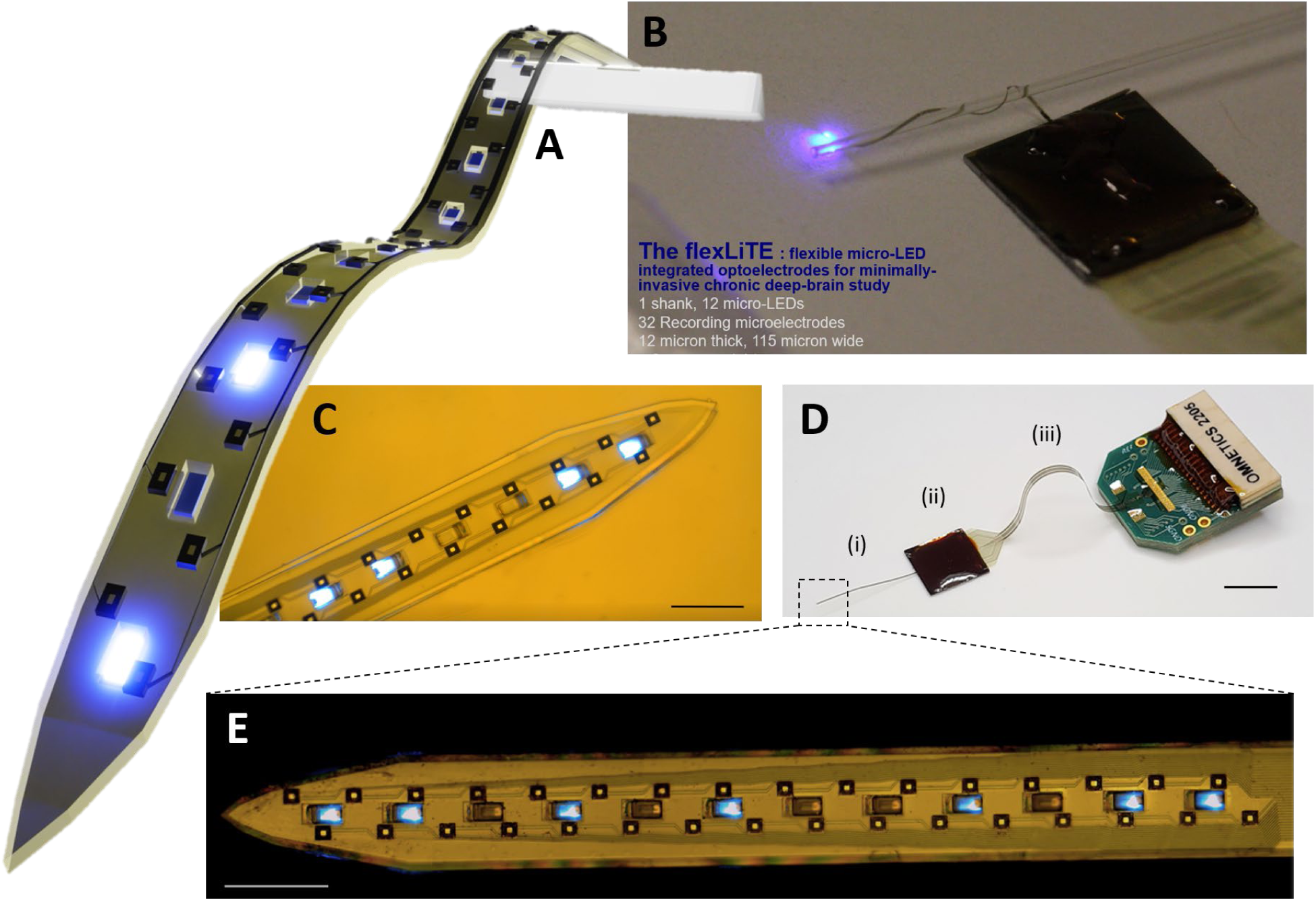
FlexLiTE optoelectrodes for chronic in-vivo experiments. **A)** Schematic image of a flexLiTE optoelectrode with 12 LEDs in a single shank with 32 recording electrodes. The backend of the probe is assembled to an interposer. **B)** A 10 mm-long shank of the flexLiTE wrapping around a 200 µm diameter optical fiber twice with LEDs turned on. Scale bar: 1 mm. **C)** Magnified image of the front end of a flexLiTE optoelectrode. Scale bar: 100 µm. **D)** Bird’s eye view of an assembled flexLiTE optoelectrode. Scale bar: 1 mm. (i) flexLiTE probe with a 1 mm long front end with LEDs and recording sites. (ii) Silicon interposer that connects the flexLiTE frontend to a cable by ball bonding. (iii) Flexible cable with metal traces for further connection to a headstage PCB. **E)** Microscope image of a flexLiTE optoelectrode with 12 LEDs and 32-recording electrodes in a single shank by stacking two modules: the LED module and the electrode module. Scale bar: 100 µm.

There are several vital reasons behind this modular approach. First, the overall yield of electronic devices is significantly affected by the number of added layers in microfabrication. The wafer fabrication yield is lowered as the number of additional mask steps increases [42–43]. This effect gets amplified when polyimide layers are added. Controlling the thickness of polymer layers is challenging when compared with other solid materials such as metal, oxide, or semiconductor due to the nature of spin coating of polymers. For this reason, we partitioned the flexLiTE optoelectrodes into two modules that are separately fabricated, tested and then assembled together. The modular approach gives significantly less burden to fabrication compared with the full monolithic integration of two modules on the same substrate, this allows for high fabrication yield. Second, the modular approach can adaptively provide a variety of configurations for different applications. For example, the relative location of recording sites and LEDs can be arranged by assembling a different combination of two modules optimized for a given application. Even extra functions can be added onto the given optoelectrode such as temperature sensing, chemical sensing, fluidic drug delivery, etc. This modular assembly integration allows for reduced manufacturing cost and time with the huge benefit of adaptability. In this work, we integrate only two modules for stimulation and recording functions through an interposer.

### Design and fabrication of µILED probe modules for optogenetic stimulation

The µILED-integrated stimulation probe module on a flexible polyimide substrate (abbreviated as ’LED probe,’ noted as ‘LED’ in Fig. 2 and Fig. 3) consists of a total of 12 micron-sized (6 µm × 15 µm) Gallium Nitride (GaN) ILEDs with a mesa height of 2.5 µm. They span ∼1 mm long top-down from the tip of the shank, and each LED can be individually controlled to stimulate neurons at cellular resolution [5].

**Fig. 2.**
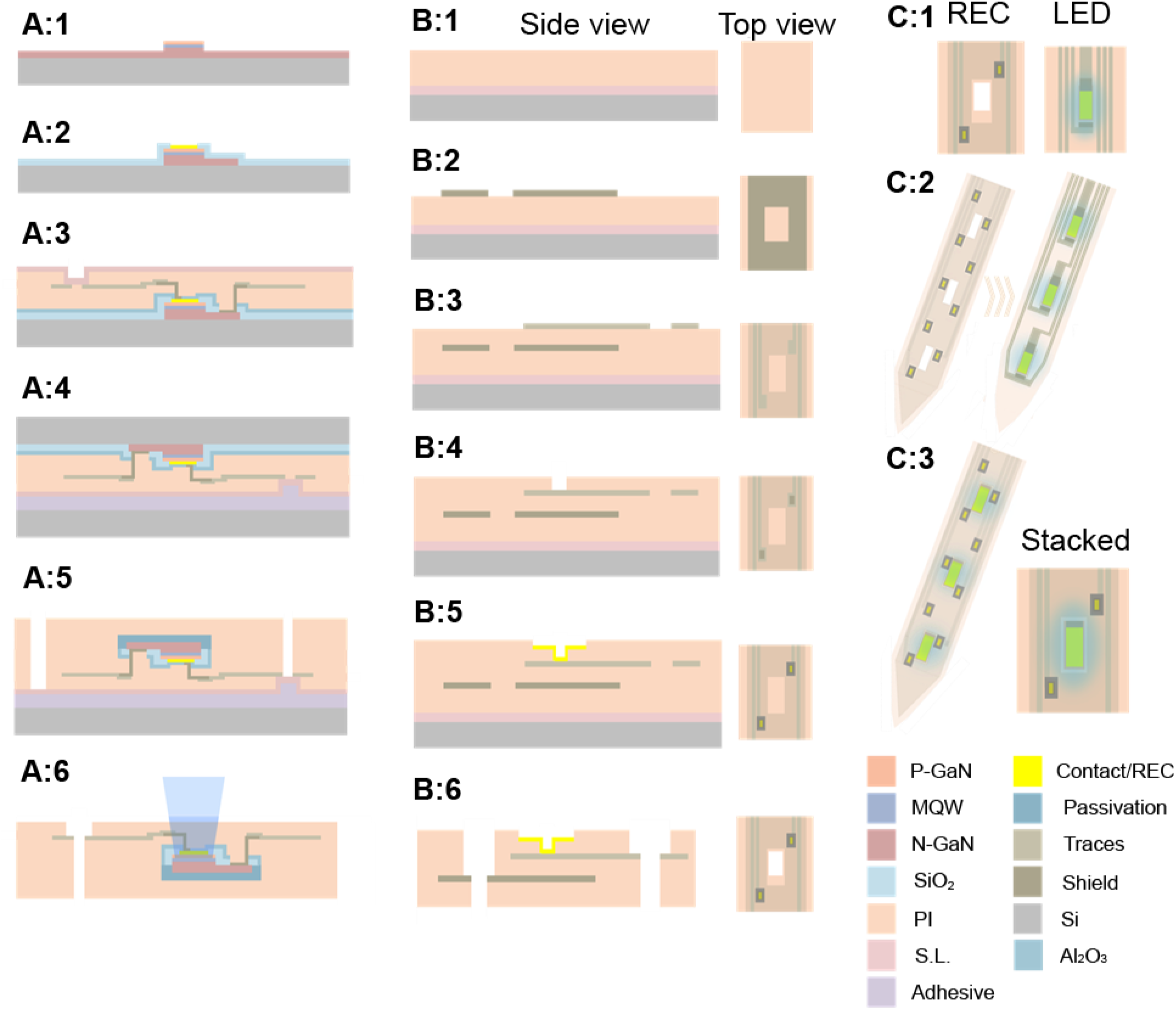
Fabrication and modular integration of flexLiTE optoelectrodes. **A)** LED flexible probe fabrication processes. **A:1)** Mesa etching of a GaN-on-silicon wafer to define μILEDs. **A:2)** Removal of n-GaN, oxide passivation and p-GaN contact metal definition. **A:3)** Additional oxide passivation, the first polyimide (PI) layer definition, contact site opening, and metal/trace definition followed by the second PI and sacrificial layer (SL) definition. **A:4)** Bonding to a transfer wafer through thermal/pressure bonding. **A:5)** Backside field layer removal LED backside reflector metal definition and the third PI passivation. Probe outline is defined, ready for release. **A:6)** Device release by SL removal. **B)** Recording electrode probe fabrication process, side view (left) and top view (right) **B:1)** the first polyimide layer definition on top of a wafer with sacrificial metal underneath. **B:2)** Metal shield defined, excluding the LED window areas. **B:3)** the second PI and metal traces for recording sites. **B:4)** the third polyimide layer and electrode via opening. **B:5)** Recording electrode definition. **B:6)** Bonding pad opening, probe outline definition, and device release. **C)** Assembly process. **C:1)** Top-view of the released REC and LED probes. **C:2)** Stacking of the REC probe on top of the LED probe. **C:3)** Stacked top view of an assembled flexLiTE optoelectrode.

**Fig. 3.**
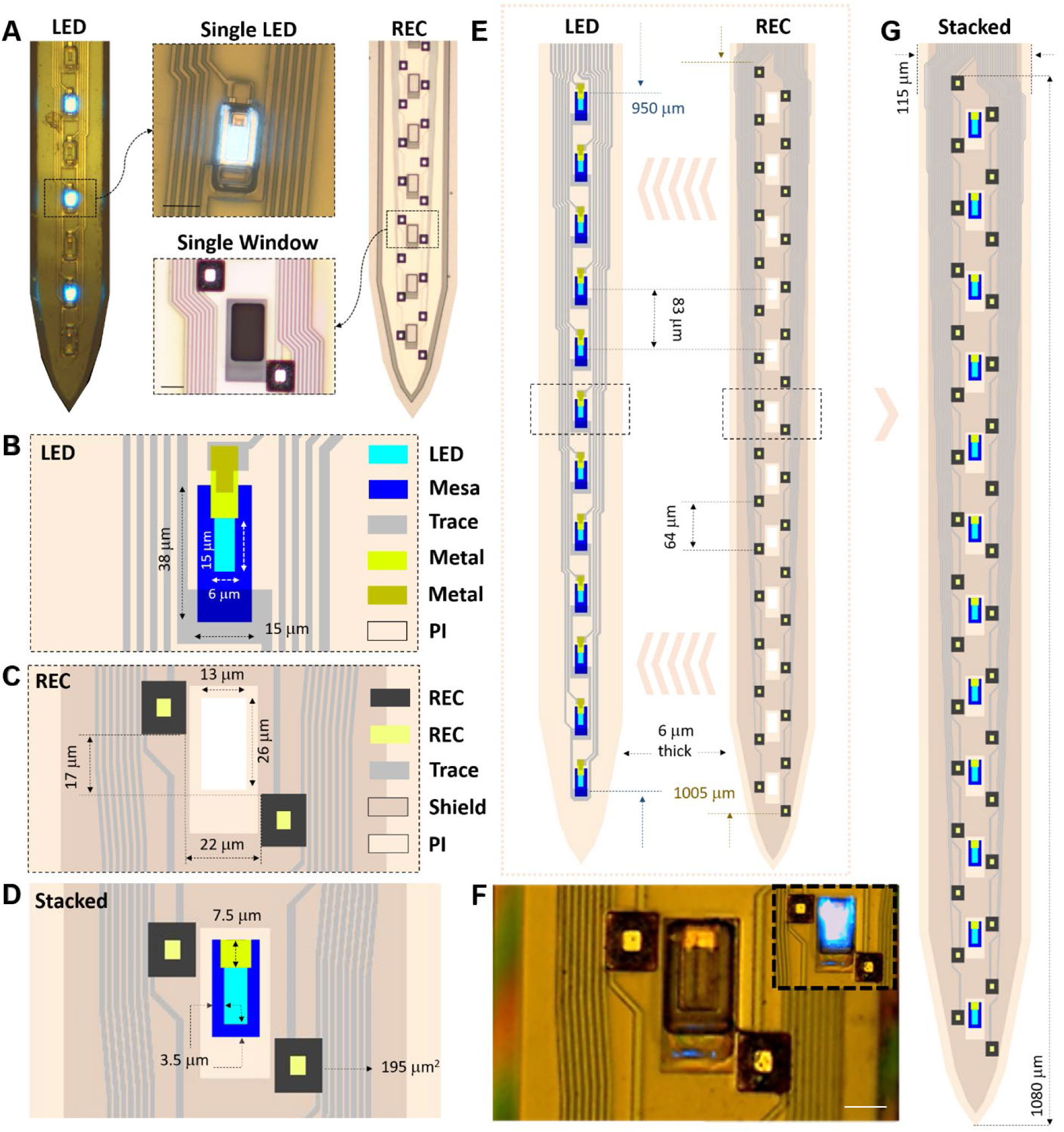
Modular stacking and assembly of flexLiTE optoelectrodes and critical dimensions. **A)** Microscope image of the released LED probe (left) and REC probe (right), with enlarged images (center) showing a single LED (top) and a single LED window and two recording electrodes (bottom) (Scale bar: 10 µm). **B)** Schematic of LEDs and dimensions. **C)** Schematic of an LED window and two recording electrodes and dimensions. **D)** Schematic of the stacked probe with an LED aligned to the window of the REC probe with alignment margins. **E)** Schematic of LED and REC probes with design parameters. The REC probe is stacked on top of the LED probe. **F)** Microscope image of the stacked probe showing a LED aligned inside the window of the REC probe. Inset shows the lighting of the LED. Scale bar: 10 µm. **G)** Schematic of the stacked probe with design parameters.

LEDs are composed of GaN quantum wells that emit blue light at λpeak ≈ 467 nm. 12 LEDs are integrated on a 6 µm-thick polyimide substrate and are connected to the backend through metal traces. The LED probe is in total 10 mm long and electrically connected to the headstage PCB through an interposer and a custom-made flexible cable (20 mm long).

The fabrication processes of the LED probe started with a GaN-based multi-quantum-well (MQW) grown on a silicon (111) wafer (4-inch, Fig. 2A:1-6). Mesa structures of LEDs were defined by two steps: one for the first mesa etching down to the n-GaN region for LED n-contacts (∼ 500 nm), and the second for isolating the mesa by etching down to the field silicon region (∼ 2 µm, a total mesa height of ∼ 2.5 µm). The silicon was exposed for the full coverage of LEDs by polyimide in the later process. A mixture of Cl2 and BCl3 gases was used for RIE definition of the mesa.

The wafer was then passivated with a thin oxide layer (∼ 500 nm) for protecting the mesa (Fig. 2A:2). The p-GaN contact was defined using Ni/Au (total 10 nm thick) metal stacks and annealed at 500 °C by rapid thermal annealing (RTA) for low contact resistance [32].

The contact holes for n-GaN were opened, followed by Al2O3 passivation. On the top the first polyimide layer (polyimide 2610, HD Microsystems, Parlin, NJ, United States) was spun and cured at 350 °C inside the oven using H2/N2 environment (thickness ∼ 1.5 µm after cure). The contact holes were etched and the n-GaN contact metal was defined along with providing p-GaN traces using the double-layer photoresist lithography for lift-off. Cr and Au layers (total 300 nm) were sputtered (Lab 18-01, Kurt J. Lesker Company, PA, US) and lifted-off. Finally, the LED p-GaN contact metal, intermediate metal traces, and n-GaN contact metal were connected to the backend ball bonding pads through 300 nm-thick metal traces of gold.

Next the second polyimide layer was spun and cured (∼ 2.5 µm). The surface of the first polyimide layer was treated with oxygen plasma for better adhesion between PI-to-PI (Fig. 2A:3). Then the bonding pads at the backend of the LED probe were opened, and the sacrificial layer (Chromium, total 225 nm) was sputtered (Lab 18-01) on the whole wafer. The wafer was transferred to a carrier wafer with an adhesive film for complete removal of the silicon substrate. The backside silicon was removed by deep-reactive-ion-etching (DRIE) (Fig. 2A:4-5). After the silicon was completely removed, the field oxide was etched away in a buffered HF solution. The exposed bottom surface of the n-GaN of LED mesa was covered by Titanium and Aluminum (200 nm).

Finally, the third polyimide layer was spun and cured (∼ 2 µm thick, ∼ 6 µm of total LED probe thickness, Fig. 2A:6). The through-holes for the ball bonding pads were opened and at the same time probe outline was defined. Then the wafer was put inside a Chromium etchant for probe release.

### Design and fabrication of REC probe modules for recording neurons

The second module of flexLiTE is the recording electrode probe (’REC probe’), which is composed of 32 electrodes. The electrodes are arranged in two rows and aligned with LEDs of the stimulation probe module (or ‘LED probe’) as shown in Fig. 1 E. Total 32 traces were defined with 1 µm metal width and 1 µm space. The LED windows were opened with a 3.5 µm misalignment margin from the edge of the LED (Fig. 2B). Fig. 3A shows the fabricated LED probe (left) and REC probe (right), and the enlarged image of the single LED illuminating through a window opening. The probe-to-probe stacking margin is illustrated in Fig. 3D with the dimensions for the probes in Fig. 3B and C. The REC probe is to be placed on top of the LED probe for close placement of the recording electrodes to the cells. The area of the recording electrode is 195 µm^2^. The vertical and lateral distances between the neighboring electrodes are 17 µm and 22 µm, respectively, while the vertical pitch of two recording electrodes in the same column is 64 µm, giving ∼ 1000 µm of covering span from top to bottom (Fig. 3E). The fabrication process flow is summarized and represented in Fig. 2B:1-6 (cross section view (left) and top view (right)).

The fabrication of REC probes started with a silicon wafer with a thin oxide grown on top (Fig. 2B:1). First, the Chromium sacrificial layer (∼ 225 nm) was defined first. Then the first polyimide layer was spun and cured (∼ 2 µm), followed by the bottom shield metal definition which works for shielding the electromagnetic interference (EMI) generated from the LED probe. The shielding layer was composed of Titanium and Gold (∼ 300 nm) and was ball bonded to the ground at the end of the assembly process (Fig. 2B:2, the side and top view of defined shield with the LED window opening).

On the top of the bottom-shield layer, the second polyimide layer was spun and cured (∼ 1.8 µm thick) as the intermediate layer between the first metal (for shield) and the second metal (for traces to connect recording electrodes and backend ball bonding pads) (Fig. 2B:3-4). The Al2O3 layer was deposited to passivate the traces (30 nm thick) [44–45]. Next, Titanium and Gold metal composites were defined for metal traces by lift-off. The top side of the traces were covered with an additional passivation layer (∼ 15 nm) and the oxide is patterned, leaving the probe-outline boundary free from any oxide covering. Next, the third polyimide layer was spun and cured (∼ 2 µm thick), followed by recording electrode formation.

The recording electrode surface was roughened to lower the impedance (Fig. 2B:5-6) [25]. Finally, the REC probe outline was defined by RIE, and devices were released in the Chromium etchant.

### Self-assembly of flexLiTE optoelectrodes using surface tension for high reproducible stacking and alignment

We introduced a thin silicon-based interposer as an intermediate assembly component to integrate the two modules: LED probe and REC probe. The interposer is composed of an array of bonding pads in the front-end for connecting the two probe modules, and another array of bonding pads in the backend for connecting a cable to the headstage PCB. Fabrication starts with a 200 µm-thick silicon wafer. A 500 nm-thick oxide layer was first deposited using plasma enhanced chemical vapor deposition (P5000 PECVD, Applied Materials Inc., CA, US). Then, metal pads and traces were defined by lifted-off. The top of the metal traces was passivated by oxide and the bonding pad openings were made. The wafer was diced into individual interposers (6 mm × 6 mm) (Fig. 1D(ii)). A cable (Fig. 1D(iii)) was prepared using similar fabrication processes to the REC probe.

The most critical assembly step is to reliably reproduce accurate alignment between the two modules. We achieve this by developing a self-aligned precision stacking technique in two-step processes. First, the LED and REC probes were stacked with coarse alignment (the REC probe on top of the LED probe) using a microscope after pretreatment with IPA. The shank tip of the stacked probe was dipped into the DI water for fine alignment. Two modules were self-aligned by surface tension (details explained in the Method section) (Fig. 2C, Fig. 3E-G). The backend of the stacked probes was ball-bonded to the interposer and then potted with epoxy. Next, the cable was ball bonded to the interposer and the headstage PCB. Finally, the Omnetics connector was attached to the PCB, and the assembled flexLiTE optoeletrodes are ready for characterization and in-vivo experiments.

### Benchtop characterization of flexLiTE optoelectrodes

In pursuance of reliable in-vivo chronic experiments, full characterization of the assembled flexLiTEs should be conducted thoroughly. Figure 4 represents the characterized results of three assembled flexLiTEs. The characterization methods are summarized in the Method section. We tested the total of 36 LEDs and 96 recording electrodes, among which 1 LED and 5 recording electrodes were not-functioning. The results represent the characteristics from 35 LEDs and 91 recording electrodes.

**Fig. 4.**
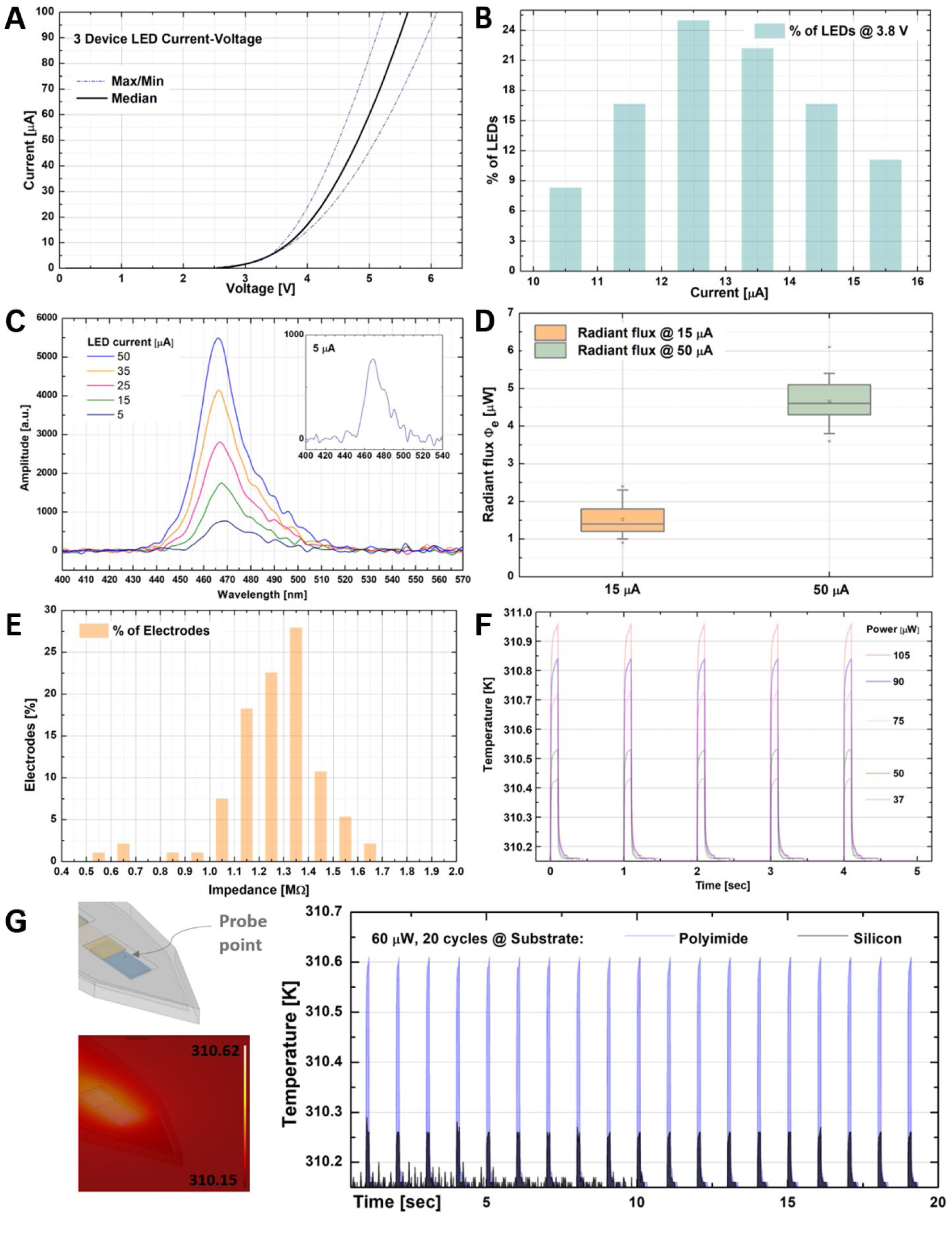
Characteristics of the fabricated flesLiTE optoelectrodes: Bench top testing from three assembled probes and thermal simulation results. **A)** LEDs current-voltage characteristics with a compliance current of 100 µA with the maximum and minimum (purple dashed line) and median (black). **B)** Distribution of LED currents at 3.8 V input voltage. **C)** LED light spectra from 5 µA to 50 µA. **D)** Radiant flux box plot for 15 and 50 µA of current. **E)** Impedance distribution of the recording electrodes measured by Intan system. **F)** FEA thermal simulation results of brain tissue heating for varied power input from 37 µW to 105 µW. Input pulse was set to the exact same square wave that was used during in-vivo testing when driving the LEDs, which is a 10 % duty cycled 1 sec long square pulse. The temperature increase was estimated in the brain tissue 8 µm above the surface of an LED. **G)** Comparison of the estimated temperature increase between polyimide optoelectrodes and silicon-shank optoelectrodes at 60 µW input power for 20 cycles.

Figure 4A shows the I-V characteristics of all LEDs measured from 0 V to 7 V under a current compliance of 100 µA. The dashed-purple lines show the maximum and minimum I-V curves from all measured results (criteria for the max/min is the voltage value at 100 µA). Compared to the LEDs formed on a silicon shank that was published in Neuron 2015 [32], the I-V curves show higher parasitic resistance due to the polyimide etching in the fabrication processes. The oxygen plasma reactive ion etching (RIE) of polyimide could leave a thin residue layer on top of the metal contact holes, leading to a degradation of the contact resistance [46]. The remedy can be adding fluorine gas in the etching process; however, this needs to be further investigated thoroughly as future work.

Figure 4B shows the percentage of the current value at 3.8 V of input voltage (Vin) for the range from 10 µA to 16 µA. Note that about ∼ 80 % of the LEDs have current values between 11 µA ∼ 15 µA (note that the Vin used during in-vivo testing was 3.8 V). The wavelength spectrum of blue-light generated from the LEDs was tested for various current levels (Fig. 4C). The inset of Fig. 4C shows the magnified view of the wave spectrum at a current level of 5 µA. The peak amplitude was measured in the range of 466-468 nm (467 nm at 15 µA of testing current).

Figure 4D shows the radiant flux (Φe) measured for 15 and 50 µA of current levels. The median values points were ∼ 1.5 and ∼ 4.5 µW for 15 and 50 µA, respectively. Considering the threshold value (optical power needed) of 1 mW/mm^2^ for activating channelrhodopsin ChR2, the optical power measured from flexLiTE optoelectrodes (∼ 1.5 µW at 15 µA) is suitable for optogenetic stimulation within 20 µm distance from the LEDs [32, 47–49].

Figure 4E denotes the distribution of impedance measured from the recording electrodes using the Intan system (see the Methods section for the test settings). More than 90 % of the electrodes showed the impedance value between 1 ∼ 1.5 MΩ, which is suitable for recording neural activities inside the target region of the brain. The impedance value was then further tracked during the in-vivo experiments over time, which will be described in the next section. As the last part of characterization, the heating effect was evaluated by COMSOL Multiphysics simulations. Low thermal conductivity of polyimide (0.12 W/m·K) may induce tissue temperature increase higher than silicon-shank µLED optoelectrodes (with the same dimension, total thickness of 12 µm) since silicon has a much higher thermal conductivity of 148 W/m·K. The 6 µm × 15 µm area (LED size, Fig. 3B) was designated as the light power generating area (blue colored region in Fig. 4G, left) in the simulation. The probing point of the temperature was set to be at 8 µm away from the LED in z-axis direction, as this would be where the neurons might be presumably located from our previous in-vivo experiments. The input power was applied for 1 sec long in a square function with a 10 % duty cycle (which is the case for our in-vivo experiments).

Figure 4F shows the corresponding temperature of the probing point during 5 cycles of various input power from 37 µW to 105 µW. These values were chosen from the actual power at the input voltage range of 3.0 V to 4.5 V. For the comparison of the polyimide vs. silicon substrate, we used the input power of 60 µW (Vin = 3.8 V, I = 16 µA) (Fig. 4G, right, total 20 cycles). The black-colored curve shown in Fig. 4G represents the temperature variation of the silicon substrate µLED probe, which is less than that of polyimide-based optoelectrodes as expected. Figures 4F and G both show that even with the continuous cycles of input power, the peak temperature value does not change, and as soon as the input pulse becomes off, the temperature rapidly goes down to the baseline temperature (which is 310.15 K = 37 °C). Even with input power of 105 µW, the difference between the temperature peak and the baseline is < 1 °C, which secures that temperature increase of the brain is negligible for in-vivo studies.

### In-vivo validation of flexLiTE optoelectrodes in a freely behaving mouse

To demonstrate the functionality of flexLITE, three fully assembled flexLiTE optoelectrodes (12 μLEDs and 32 recording electrodes) were chronically implanted into the dorsal hippocampus (AP: -1.34mm, ML: -1.00mm and DV: -2.3mm) of a Tg Thy1-ChR2/EYFP mouse. Each flexLiTE was inserted using a glass pipette as a mechanical shuttle (detailed description in the Methods section, Fig. 5A and B). After reaching the target depth, the glass pipette was retracted leaving the flexLiTE probe inside the brain. Opto-electrophysiology recordings from freely moving mice started 1 week after the surgery from freely moving mice (Fig. 5-6, Fig. S1).

**Fig. 5.**
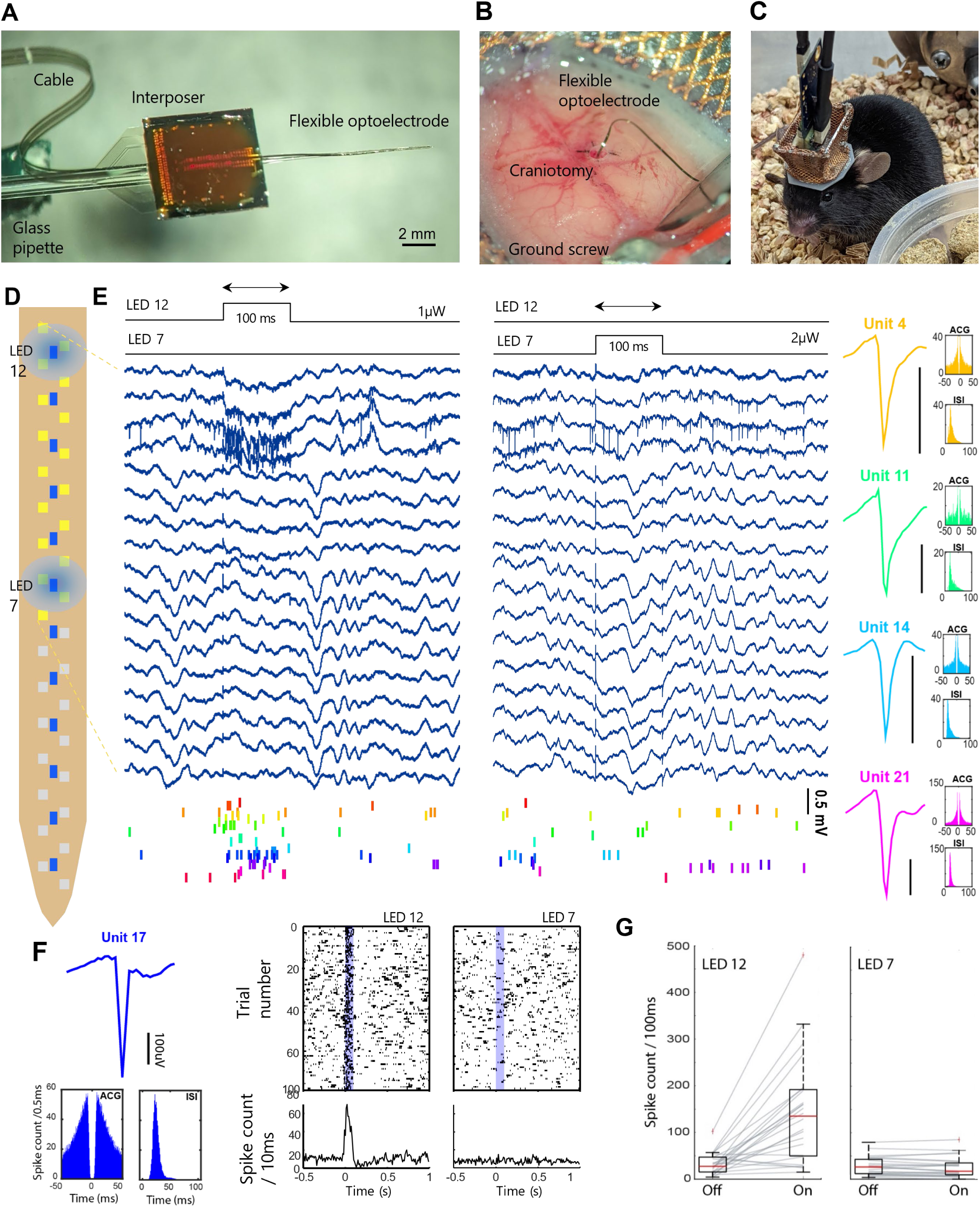
In vivo opto-electrophysiology with flexLiTE optoelectrodes in freely moving mice: short-term device functionality demonstration. A-B) Implantation of flexLiTE optoelectrodes. **A)** A flexLiTE optoelectrode is attached to a glass pipette using polyethylene glycol. **B)** After insertion, the glass pipette is retracted leaving behind the flexiLiTE. **C)** Image of a mouse during experiment with the headstage and micro-LED driving cable connected to PCB connectors of flexLiTE. **D)** Schematic of the flexLiTE configuration. Middle column (blue squares): location of 12 µILEDs; left and right column (yellow and grey squares): location of 32 recording sites (recorded signals from yellow-colored recording channels are shown in E). (**E)** Respective data collected from recording channels mapped yellow color coded in (D) to demonstrate the functionality of the flexLiTE within different locations. The experiment was held after 2 weeks from implantation. The spiking data was quantified for 24 isolated cells (22 putative pyramidal and 2 putative interneurons) recorded simultaneously from CA1. Represented signals are traces of the recorded wide-band signals (0.1-7500 Hz) and the raster plots of sorted units when LEDs 12 (left panel) and 7 (middle panel) were triggered. Mean waveform of 4 representative cells and their corresponding ACG and ISI histograms are shown in the rightmost panel (black bar represents 100 µV). Location of LED # 12 and 7 are in (D). LED # 12 and 7 were driven with 1 µW and 2 µW, respectively. **F)** Peristimulus time histogram for Unit # 17 during 100 light pulses (1 sec pulse, 10 % duty cycle with 1 µW power) for LED # 12 and 7, respectively. **G)** Spike count / 100 ms where LED # 12 and 7 was individually selected and pulsed.

**Fig. 6.**
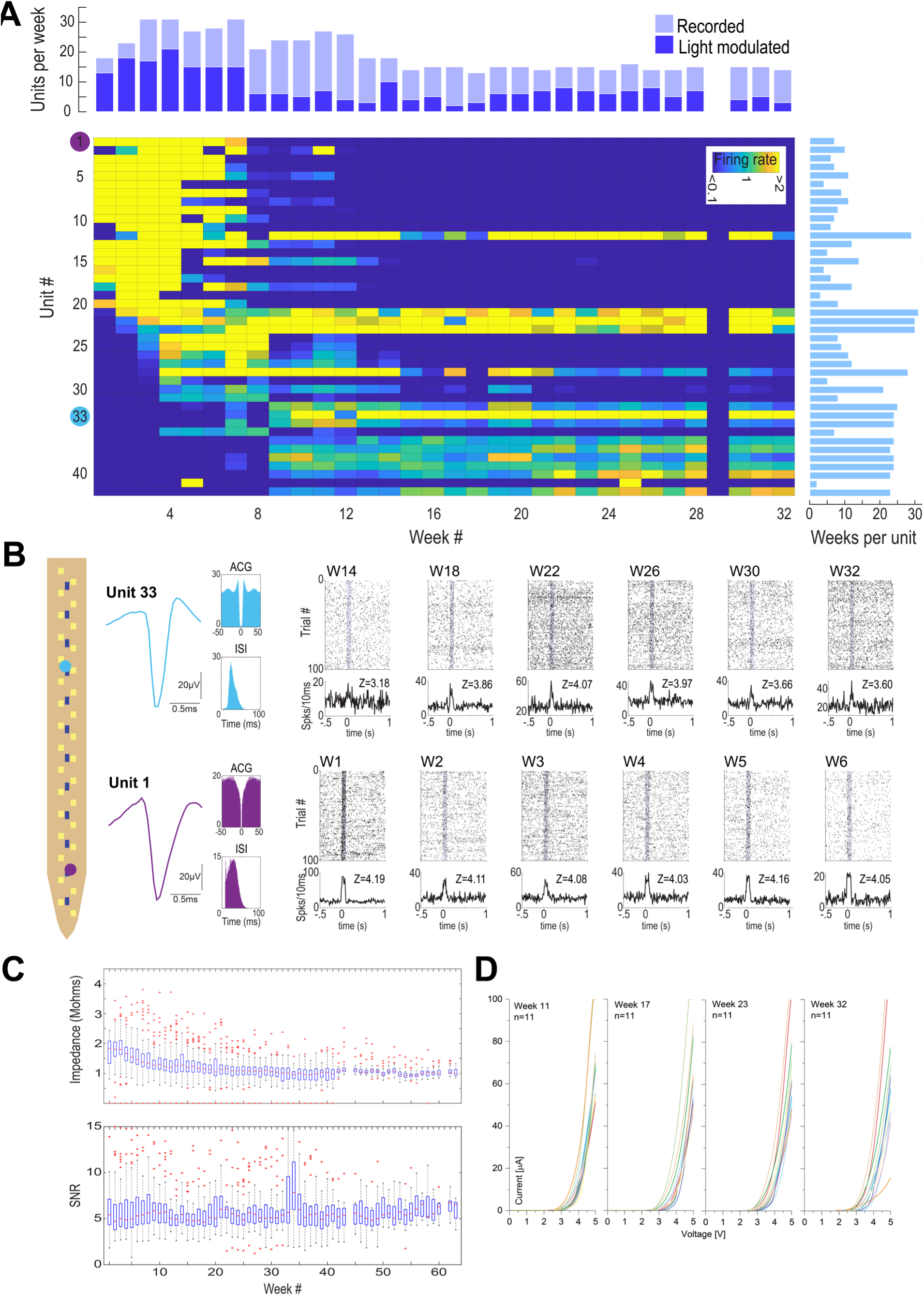
In vivo opto-electrophysiology with flexLiTE optoelectrodes in freely moving mice: Device lifetime validation through 8-month chronic experiment. **(A)** Heatmap shows the color-coded firing rate of all 42 recorded cells throughout 32 weeks from device 2. The number of recorded cells and the number of cells positively modulated through blue light is presented from week 1 to 32 (top). The number of weeks per cell is shown to the right. **(B)** Schematic of the flexLiTE with 12 micro-LEDs (blue) and 32 recording electrodes (yellow) with the location of the example cell # 1 and 33 (left). Autocorrelation histogram and PSTH for 100 stimulation trials with Z score for unit 33 and 1 as a representative example for long-term (#33) and short-term (#1) recorded cell. **(C)** Weekly impedance and signal to noise ratio for 3 chronically implanted devices (device 1 = 32 weeks, device 2 = 63 weeks and device 3 = 41 weeks). **(D)** Device 2 micro-LED current-voltage characteristics for V = 0 to 5 V vs. I = 0 to 100 µA for week 11, 17, 23, and 32 after implantation.

For a short-term validation, we recorded light-induced neural responses from one of the flexLITE (device 1) at week 2. A cell was classified as “recorded” when its firing rate during the specified week’s baseline was above 0.1 Hz and labeled as “light modulated” when its firing rate during the illumination period of any micro-LEDs significantly increased when compared to the off period (detailed description of the analysis can be found on the Methods section). During week 2, device 1 recorded a total of 24 single units, 22 putative pyramidal neurons and 2 putative interneurons (Fig. 5E-F, the details of cell classification criteria can be found in the Method section). Half-second full-band peristimulus recordings, single unit waveforms and raster plots are shown in Fig. 5E. Left panel corresponds to the recorded signals when LED # 12 is illuminated (1 sec long pulse with a 10 % duty cycle, LED optical power was set to 1 µW), while right panel shows when LED # 7 is turned on-and-off (1 sec long pulse with a 10 % duty cycle @ 2 µW). Activation of LED #12 evokes a significant increase in the spiking activity in the neighboring neurons (Fig. 5E-G). In contrast, LED #7 did not evoke a clear response in the same neurons regardless of using doubled power (Fig. 5E-G). Figure 5F shows the waveform, auto-correlogram and inter-spike interval, as well as the resulting raster plots and peristimulus time histograms (PSTH) of an example putative pyramidal neuron (unit 17) recorded during the session. The total number of spikes for each unit during the illumination period is shown in Figure 5G and compared to the rolling average of an identical off-period for both LEDs (7 and 12), further confirming the local effect of a single LED.

To demonstrate the long-term chronic functionality, weekly recordings were performed throughout the lifespan of all 3 devices (Table 1, Fig. 6). A single session consisted of 15 minutes of baseline and immediately after stimulation trains for the selected micro-LEDs were given. A stimulation train consisted of 100 cycles of 100ms ON and 1900ms OFF.

**Table 1.** Chronic In-vivo experiment – long term demonstration.

| Chronic | # of functional electrodes /32 | # of functional LEDs /12 | Light modulation + electrophysiology Longevity*** | Device LED Longevity (failure mode) | Device REC Longevity (failure mode) | Z \| week1 Post implant | Z \| week @ fail time Post implant | LED I @ V = 4.5 V week3 | LED I @ V = 4.5 V @ *** |
| --- | --- | --- | --- | --- | --- | --- | --- | --- | --- |
| Device 1 | 28 | 12 | Week 12 | Week 16 (shorted) | Week 39 (signal lost) | $2.00 \pm 0.22 \text{ M}\Omega$ | $1.20 \pm 0.14 \text{ M}\Omega$ | $36.11 \pm 5.22 \text{ }\mu\text{A}$ | $31.325 \pm 8.10 \text{ }\mu\text{A}$ |
| Device 2 | 31 | 11 | Week 32 | > Week 57 (explanted) | Week 40 (signal lost) | $1.73 \pm 0.48 \text{ M}\Omega$ | $0.92 \pm 0.20 \text{ M}\Omega$ | $40.29 \pm 13.72 \text{ }\mu\text{A}$ | $39.25 \pm 22.08 \text{ }\mu\text{A}$ |
| Device 3 | 32 | 11 | Week 16 | Week 25 (shorted) | Week 59 (signal lost) | $1.57 \pm 0.25 \text{ M}\Omega$ | $1.28 \pm 0.16 \text{ M}\Omega$ | $30.72 \pm 7.39 \text{ }\mu\text{A}$ | $35 \pm 15.04 \text{ }\mu\text{A}$ |

After concatenating weekly recordings for all 3 devices, the average recording longevity was 45.33 ± 15.95 weeks and the average longevity for the micro-LED module was 32.67 ± 21.55 weeks (details given in Table 1). A total of 130 single units were sorted from all 3 recordings. In average, a total of 43.33 ± 6.11 individual cells were sorted throughout the recording lifespan of each device. The averaged weekly single unit yield from each device was 15.28 ± 6.77, with a peak of ∼29 single units per device at weeks 3 and 4. For the 130 single units recorded from all 3 devices, a single unit was recorded in average for 14.06 ± 13.25, with a maximum of 59 weeks; and 112 out of the 130 recorded neurons were detected for ≥ 2 weeks. In average a total of 9.86 ± 5.98 neurons per week were positively light-modulated, with a maximum average during the first 4 weeks of 16.25 ± 4.66.

Long-term tracking of a second device (device 2) is shown as an example in Fig. 6 A-B. The firing rate for each sorted cell throughout the duration of the device is shown in Fig. 6A. On average, we recorded a total of 20.81 ± 6.82 cells per week, out of which 8 ± 4.99 were positively light-modulated during the stimulation period. Figure 6A also shows the number of weeks each cell was recorded, the longest reported was 30 weeks for this device. On average, cells were recorded for 14.16 ± 9.1 weeks. Figure 6B shows the schematic of the flexLiTE with the corresponding position of 2 representative recorded cells in the middle and tip-side. Depending on the position of the group of cells, three of the 12 micro-LEDs were used for stimulation/recording functionality demonstration (LED # 2, 8, and 10; note that LED # 1 is the one closest to the tip, Fig. 6B). The trilaterated position for 2 representative neurons is shown in Figure 6B (units 1 and 33, blue- and purple-color circles in Fig. 6A). These examples display clear modulation in response to the activation of the nearest micro-LED according to their peristimulus time histogram (PSTH), raster plot and the resulting Z-score (Raster plots and PSTHs for every neuron and week are shown in Fig. S1).

Figure 6C shows the impedance change of the recording electrodes over time from all 3 devices. The impedance median value from all 3 devices throughout their lifespan was 1.12 MΩ, with a maximum median range between ∼1.8 and ∼1.3 MΩ during the first 8 weeks after implant and a median signal-noise-ratio of 5.28 (Fig. 6C). Fig. 6D shows the variation in the I-V characteristics of the micro-LEDs throughout 32 weeks for the best performing device, where all working 11 LEDs did not show sign of failure (open or short) when tested at the specified periods. In this case, the micro-LED module outlasted the electrophysiology module despite presenting nominal impedance values, spiking signal was completely lost and no spontaneous nor evoked single units were detected (Table 1). For this device, the long-term device lifetime was characterized through the impedance measurements and micro-LEDs I-V curves. The device presented no shorted/opened LEDs throughout the experiment (13 months Fig. S2 and S3). Altogether, these results demonstrate the functionality and reliability of the flexLiTE optoelectrode.

## Discussion

The flexLiTE optoelectrode was demonstrated for the first time for chronically modulating and recording the brain neuronal circuitry through chronic in-vivo experiments for up to 8 months after implant. The modular structure of flexLiTE can expedite the neuroscience study in terms of both efficiency in surgery and device assembly yields. The modular design (‘LED probe’ and ‘REC probe’) can be separately fabricated and assembled on one substrate via an interposer. This enhances the yield of probes and provides fast turn-around time since each module can be selectively assembled depending on the end users’ needs, simplifying manufacturing processes and furnishing versatility. This modular reconfigurable approach allows the probe backend to be elongated into any length (in this work the length of the cable was 20 mm but it can be elongated at any preferred length) to meet the need that the neuroscience experiment requires. The main design for the prototype probe was to cover a 1 mm span from a target region, in this case the dorsal hippocampus, with 12 LEDs and 32 recording electrodes, respectively, on an ultra-flexible polyimide substrate for long-term chronic experiments. These device elements were incorporated onto a polyimide substrate with a total dimension of 10 mm long (only the shank region), 115 µm of shank width, and 12 µm of shank thickness. By virtue of matched surface energy of polyimide, we could realize multi-module-stacked structures where the REC probe was placed on top of the LED probe with precise alignment from surface tension (< 3.5 µm tolerance) when placed and dried up in an aqueous solution such as IPA. The characteristics of the LEDs and recording electrodes, including the thermal simulation results of the brain tissue heating due to LED driving, showed that flexLiTE optoelectrode is suitable for in-vivo experiments. FlexLiTE optoelectrodes were implanted in the CA1 region of the hippocampus of the mouse. Once the tissue was healed after two weeks of implantation, chronic in-vivo experiments were conducted to validate the capacity of the flexLiTE as a fully functional chronic optoelectrode. First, the REC probe functionality was demonstrated with single unit recordings. Secondly, the optogenetic neuromodulation was validated by recording of LED illumination-induced spikes at 467 nm in wavelength (blue light). Compared with non-illuminating states we revealed the stimulation-induced spiking rates increased significantly.

By selectively triggering specific LEDs we demonstrated the high spatiotemporal capabilities of the flexLITE optoelectrodes for delivering light and local modulation of the neuronal population. Lastly, the chronic-capability was validated by conducting the same in-vivo experiments over time. The spiking rate discrepancy between non-/ and illuminating states demonstrated that the firing rate of the recorded cells was modulated significantly up to 32 weeks Our modular architecture of assembly is easily scalable to increase the density of μLEDs and recording electrodes by design variation, making flexLiTE optoelectrodes suitable for large-area chronic monitoring of neural circuits and targeted neuromodulation in long-term behavioral studies. In conclusion, flexLiTE devices are suitable for longitudinal studies, enabling neuroscientists for the first time to perform reliable electrophysiological recordings paired with optogenetics for extended periods.

## Methods

### flexLiTE fabrication and assembly

The LED probes, REC probes, interposers and cables were fabricated in the Lurie Nanofabrication Facility at the University of Michigan, Ann Arbor, Michigan. GaN-on-silicon wafers were purchased from Enkris (Enkris Semiconductor Inc., Suzhou, China). Polyimide 2610 and 2611 were spun and cured by the Laurell Spinner (Model Ws-650-23B, PA, United States) and Vacuum Oven (YES PB8-2B-CP, Yield Engineering Systems, CA, United States), respectively. The LED and REC probe traces were passivated for longevity enhancement by coating of Al2O3 (Oxford OpAL ALD). At the end of the front-side fabrication the wafer was coated with a PI adhesive and bonded to a carrier wafer by a wafer bonder (EVG 510, EV group, Sankt Florian am Inn, Austria). Two-step probe outline definition incorporates a mid-temperature curing for 1 hour, patterning through plasma RIE (Plasmatherm 790, Plasma-Therm, LLC, FL, United States), and high-temperature curing for 1 hour before released in a Cr etchant (Chromium Etch 1020, Transene Company, Inc., MA, United States). Polyimide-to-polyimide interfaces were surface treated with oxygen plasma (Tergeo Plasma cleaner, PIE Scientific LLC, CA, United States). Interposers were fabricated using a 200 µm-thick silicon wafer and diced into 6 mm × 6 mm squares before assembly. All the lithography for the LED/REC probes was done using a stepper (GCA AS200 AutoStep) and the interposers and cables were patterned by a contact aligner (MA/BA-6 Mask/Bond Aligner).

Assembly processes except for the PCB fabrication (PCBWay, Shenzhen, China) was done in-house (EECS building, University of Michigan, Ann Arbor, MI, United States). Before the assembly, individually manufactured LED/REC probes were first surface treated with IPA (Isopropyl alcohol) making surfaces hydrophilic, and two probes are stacked (REC probe on top of the LED probe) for coarse alignment. Next, the probe shanks of the stacked probes were dipped and pulled out slowly from the DI water container where fine adjustment of self-alignment took place from surface tension. Next, the stacked probes were ball bonded (K&S 4524-D, Kulicke and Soffa Industries, Inc., PA, USA) to the interposer using aluminum bumps. The 8 μm-thick, 20 mm-long custom-made polyimide (PI2611) cables connected the interposer to the headstage PCB by ball bonding. The 36-pin and 18-pin connectors (Omnetics Connector Corporation, MN, Unites States) were soldered to the PCB. Electrical connections of the fully assembled device were passivated by Epoxy (EPO-TEK 353ND-T, Epoxy Technologies, MA, USA).

### Electrical and optical characterization

After flexLiTE probes were assembled, we characterized the current vs. voltage (I-V) and radiant flux vs. current (Φ*e*-I) for LEDs and the impedance measurement for the recording electrodes (Ω-channel #). A total of 3 probes were used for bench-top characterization. The I-V characteristics were measured using the source-meter (Keithly 2400, OH, US) and the Φ*e*-I was measured using an optical power meter composed of an integrating sphere and spectrometer (FOIS-1, and Flame, Ocean Optics, FL, US). The entire probe (10 mm-long probe shank part) was placed under an integrating sphere. The size of the interposer was small enough to put the flexible probe inside the sphere to ensure the light from the LEDs are delivered to the optical measuring system. Optical power integration range was set to 400 – 500 nm in wavelength, where the maximum power was measured at 467 nm of wavelength. The recording site impedance was measured at 1 kHz by Intan system (Intan RHD 32-channel recording headstage and RHD2000 USB interface board, Intan Technologies, CA, US). The probe shank was placed in a 1× phosphate buffered saline solution (MP Biomedicals, OH, US) and the impedance was measured for all 32 channels.

### Thermal simulation

The flexLiTE LED design and configuration was taken into the COMSOL Multiphysics for thermal simulation. The same dimension of the device fabrication mask layout was used. The temperature increase was estimated in the tissue 8 µm above the LED surface. (Note that the thickness of the REC probe is ∼ 6 µm, and the second PI layer for the LED probe is ∼ 2 µm.) The probe shank was presumed to be surrounded by brain tissue. The actual electrical power was applied as input power for the simulation from a voltage range of 3.0 V to 4.5 V. For simplicity of thermal modeling, we assumed that the LED probes were composed of only polyimide material with no metal traces for the worst-case simulation. The bio-heat transfer equations were used from [50]. All the related parameters were taken from [32] except for polyimide. We used a thermal conductivity of 0.12 W/m·K for the polyimide in the COMSOL simulation.

### Animal experiments: In-vivo opto-electrophysiology

All experiments were approved by the Institutional Animal Care and Use Committee at New York University Medical Center. Animals were handled daily and accommodated to the experimenter before the surgery and homecage recordings. Three adult male mice (Thy1-ChR2, ∼31 g) were kept in a vivarium on a 12-hour light/dark cycle and housed up to 3 per cage before surgery and individually after it. Atropine (0.05 mg kg^−1^, s.c.) was administered after isoflurane anesthesia induction to reduce saliva production. The body temperature was monitored and kept constant at 36–37 °C with a DC temperature controller (TCAT-LV; Physitemp, Clifton, NJ, USA). Stages of anesthesia were maintained by confirming the lack of a nociceptive reflex. The skin of the head was shaved, and the surface of the skull was cleaned by hydrogen peroxide (2%). A custom 3D-printed base plate (Form3 printer, FormLabs, Sommerville, MA) [51] was attached to the skull using C&B Metabond dental cement (Parkell, Edgewood, NY). A stainless-steel ground screw was placed above the cerebellum and sealed with dental cement (Unifast LC, GC America). The location of the craniotomy was marked (AP: -1.34mm, ML: -1.00mm) and a 700-µm craniotomy was drilled. After the dura was removed the flexiLiTE optoelectrode was inserted into the brain (DV: - 2.3mm from the surface of the brain) using a glass pipette with a tip diameter of 15-20 µm (3-000-203-G/X, Drummond Scientific, PA). The glass pipette was retracted, and the craniotomy was sealed with dura-gel (Cambridge Neurotech) and then covered with dental cement. Finally, a protective cap was built using copper mesh [54]. The mouse recovered for at least 7 days after surgery. Animal was recorded in its homecage. The collected data was digitized at 20 kS/s using an RHD2000 recording system (Intan technologies, Los Angeles, CA).

### Animal experiments: Controlling flexible µLEDs

Current-controlled stimulation was used to drive individual μLEDs (OSC1Lite [39]), 12-ch current source (https://github.com/YoonGroupUmich/osc1lite). The stimulation amplitude and waveform were defined using OSC1Lite’s open-source graphical user interface.

### Animal experiments: Single unit analysis

A concatenated signal file was prepared by merging all recordings from each week for each subject. Putative single units were first sorted using Kilosort [52] and then manually curated using Phy2 and custom plug-ins (https://phy-contrib.readthedocs.io/, https://github.com/petersenpeter/phy2-plugins). After extracting timestamps of each putative single unit activity, peristimulus time histograms and firing rate gains were analyzed using a custom MATLAB (Mathworks, Natick, MA) script. To measure the effect of light on spiking, peristimulus time histograms (PSTHs) were built around stimulus onset (spike trains were binned into 10-ms bins). Baseline and light-induced spiking rate were calculated for each single unit. Baseline was defined as light-free epochs (1900 ms) between trials and stimulation period as µLED was on (100 ms). A Neuron was classified as “optically modulated” when the z-score corresponding to the ON period was ≥ 2. Z-score was calculated for 100ms-bins PSTH. The signal-to-noise ratio for each cell was calculated weekly by dividing the averaged peak amplitude from the largest amplitude channel by the standard deviation of the band-passed (500-8000Hz) recording from the same channel.

### Animal experiments: Cell-type classification

Recorded cells are classified into three putative cell types: narrow interneurons, wide interneurons, and pyramidal cells. Interneurons are selected by 2 separate criteria. Narrow interneuron is assigned if the waveform trough-to-peak latency is less than 0.425 ms. Wide interneuron is assigned if the waveform trough-to-peak latency is more than 0.425 ms and the rise time of the autocorrelation histogram is more than 6 ms. The remaining cells are assigned as pyramidal cells. Autocorrelation histograms are fitted with a triple exponential equation to supplement the classical, waveform feature based single unit classification (https://cellexplorer.org/pipeline/cell-type-classification/) [53]. Bursts were defined as groups of spikes with interspike intervals of < 9 ms.

### Animal experiments: In-vivo impedance measurement

The impedance of the recording probe was measured during each recording session (n=5 sessions, Fig. 6) using an RHD USB interface board (Intan Technologies LLC, CA, USA). The impedance test measurements were performed at 1 kHz frequency. This measurement provides a quick and rough estimate of the quality of the recording sites.

## Author contributions

E.K. and J.R.L.R contributed equally to this manuscript. E.K. and E.Y. defined the conceptual design of flexLiTE. E.K. designed, fabricated, and assembled flexLiTE optoelectrodes; conducted CAD simulations; and conducted benchtop tests. E.K. and M.-

L.H. characterized the flexLiTE optoelectrodes. J.R.L.R. and M.V. conducted the in-vivo experiments and processed/analyzed the recorded neural signals. G.B. and E.Y. supervised the study. E.K, J.R.L.R, and M.V. wrote the manuscript. All authors discussed the results and commented and edited the manuscript.

