## Supplementary Information for "flexLiTE: flexible micro-LED integrated optoelectrodes for long-term chronic deep-brain studies"

**Supplement Figure 1**. Peristimulus histogram for all recorded cells from week 1 to week 32 (week 29 was skipped) from one device with cell unit number noted in each.

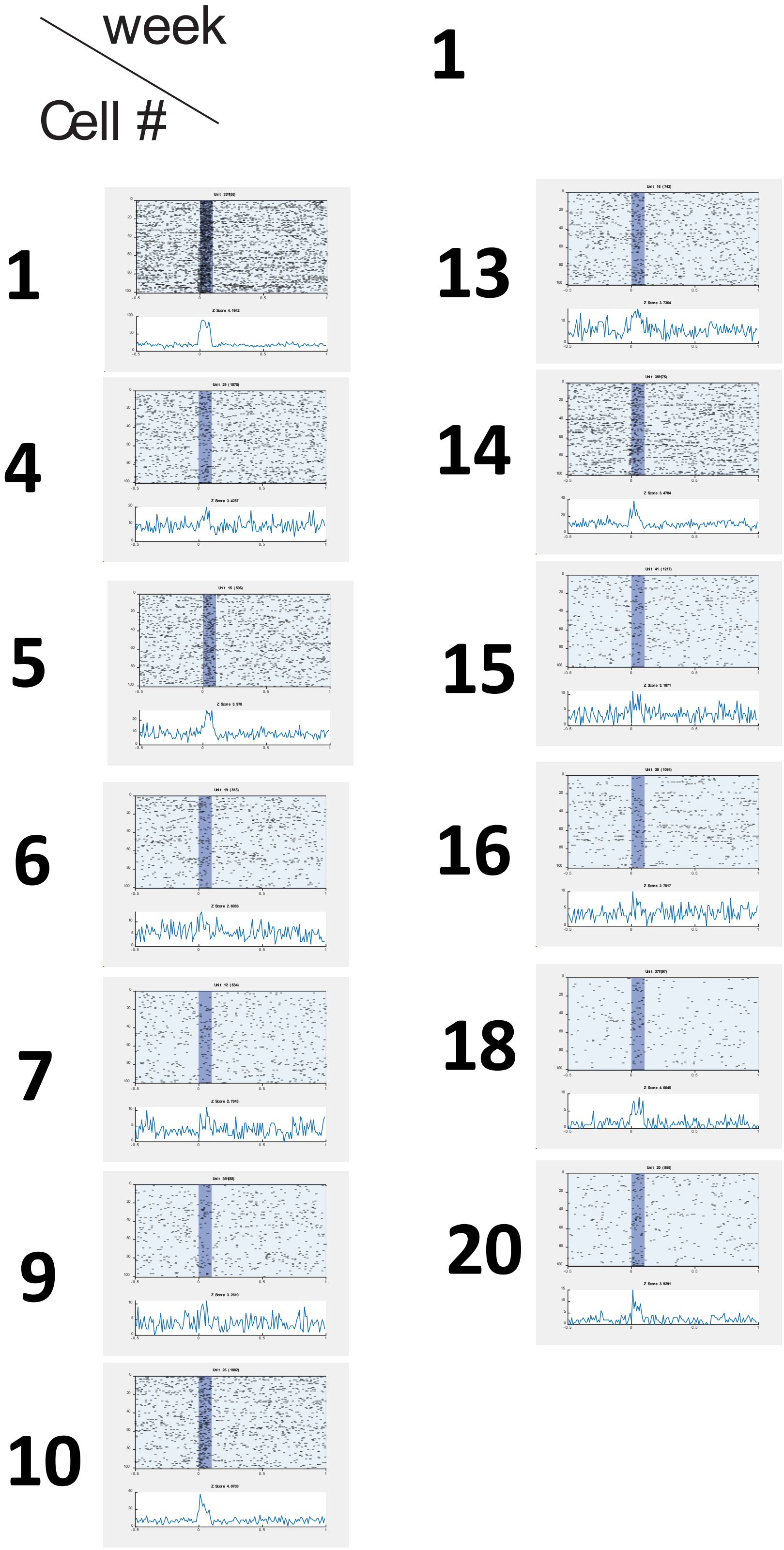

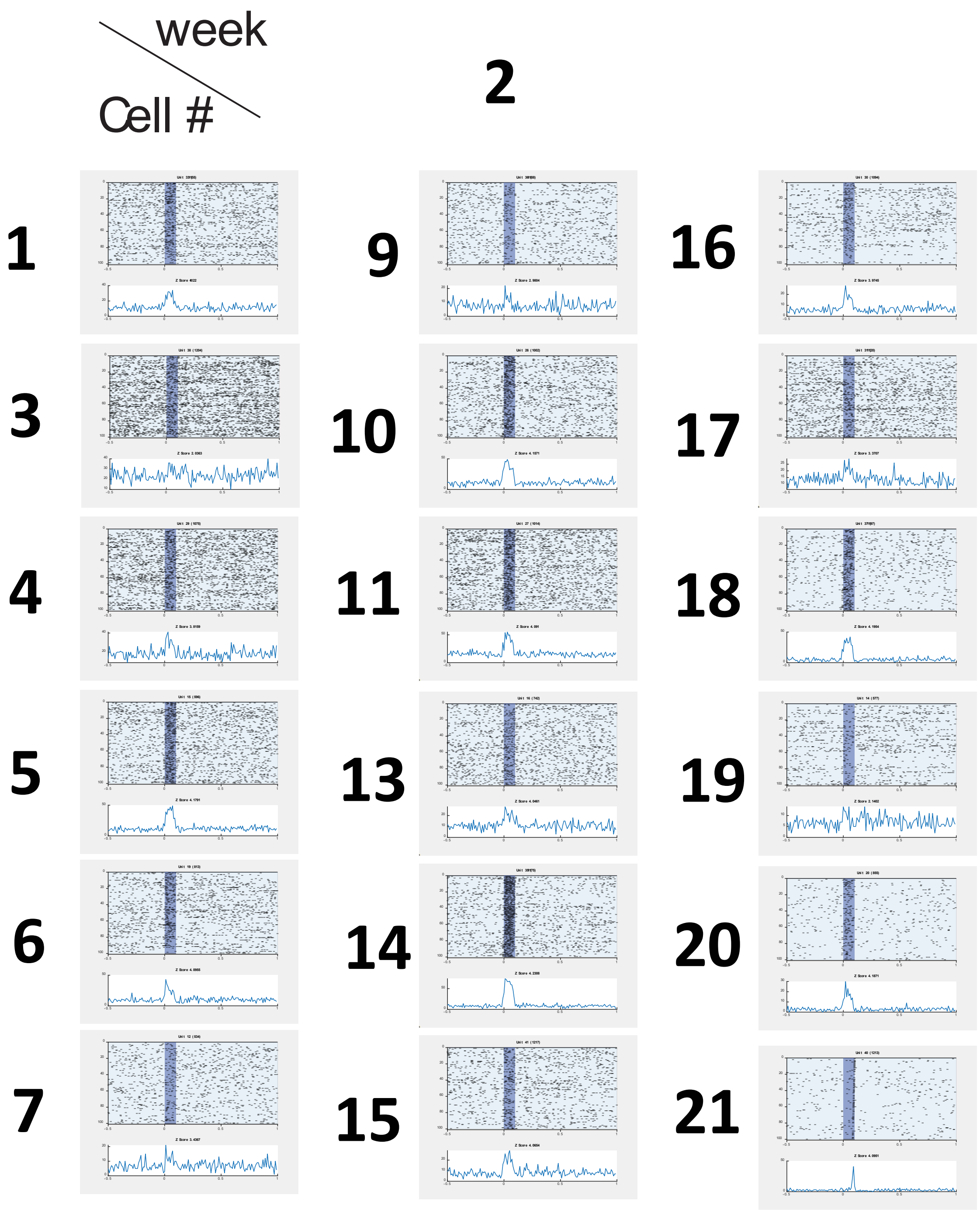
week
Cell #
2
1
9
16
3
10
17
4
11
18
5
13
19
6
14
20
7
15
21

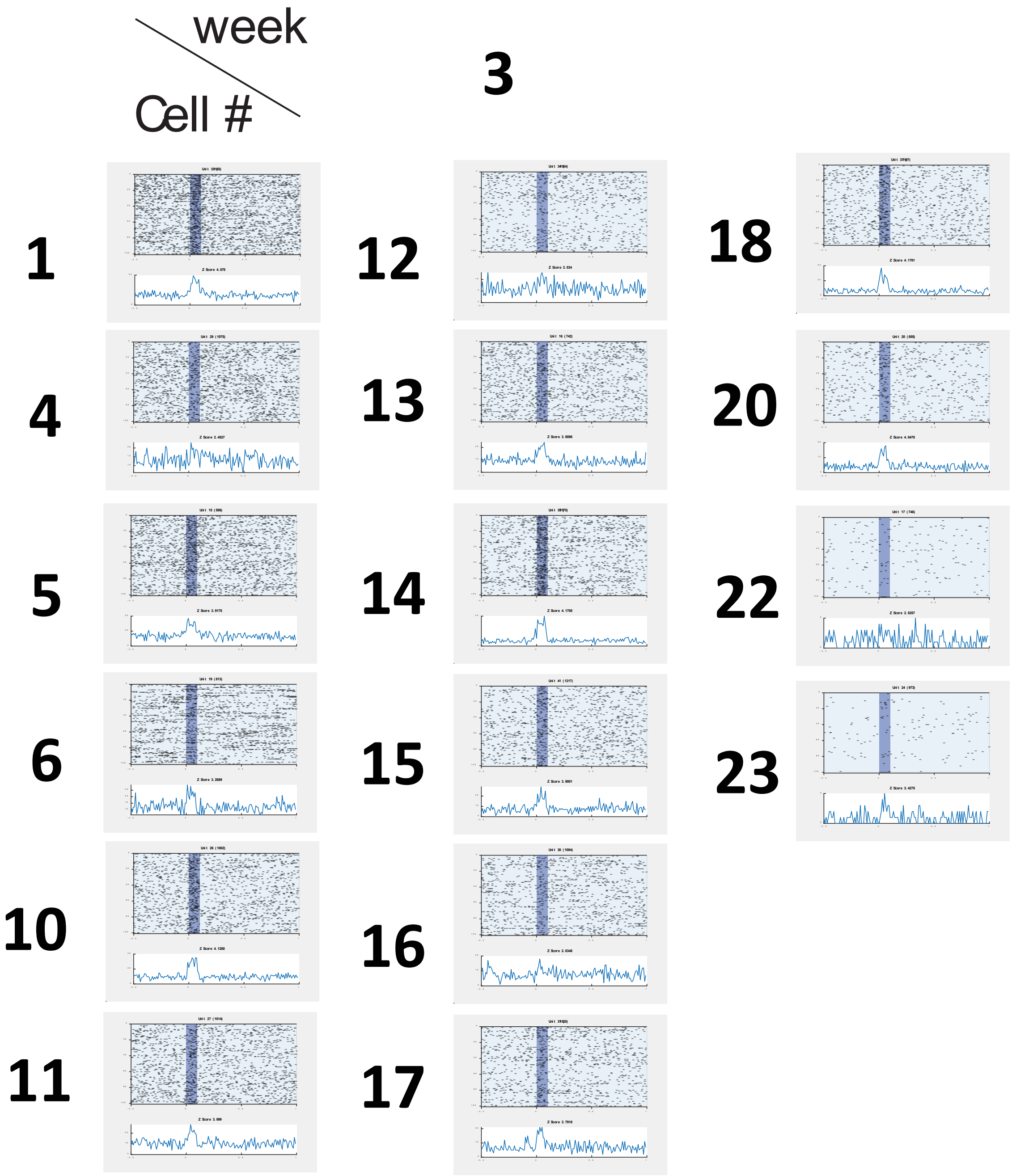
week
Cell #
3
1
12
18
4
13
20
5
14
22
6
15
23
10
16
11
17

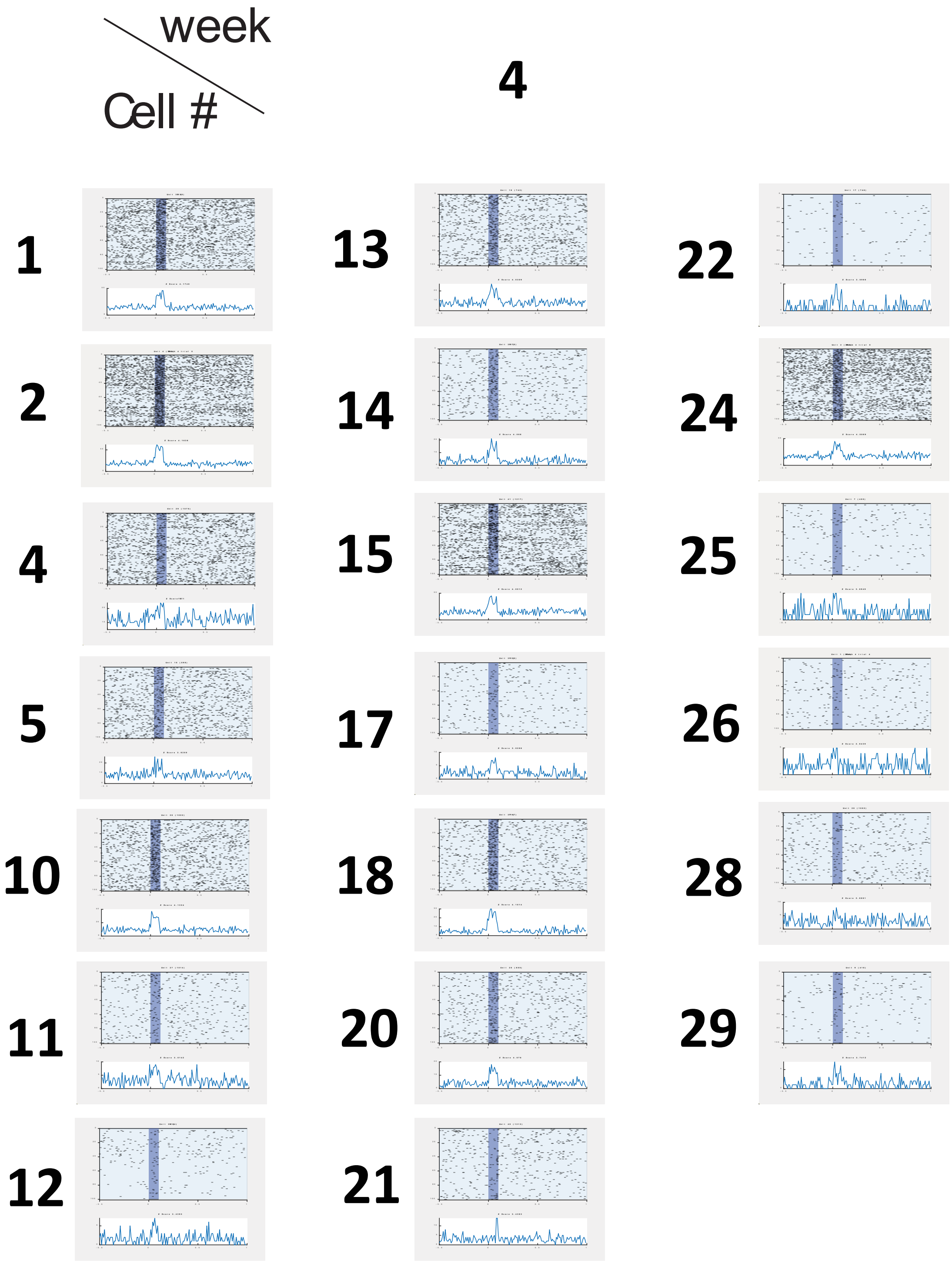
week
Cell #
4
1
2
4
5
10
11
12
13
14
15
17
18
20
21
22
24
25
26
28
29

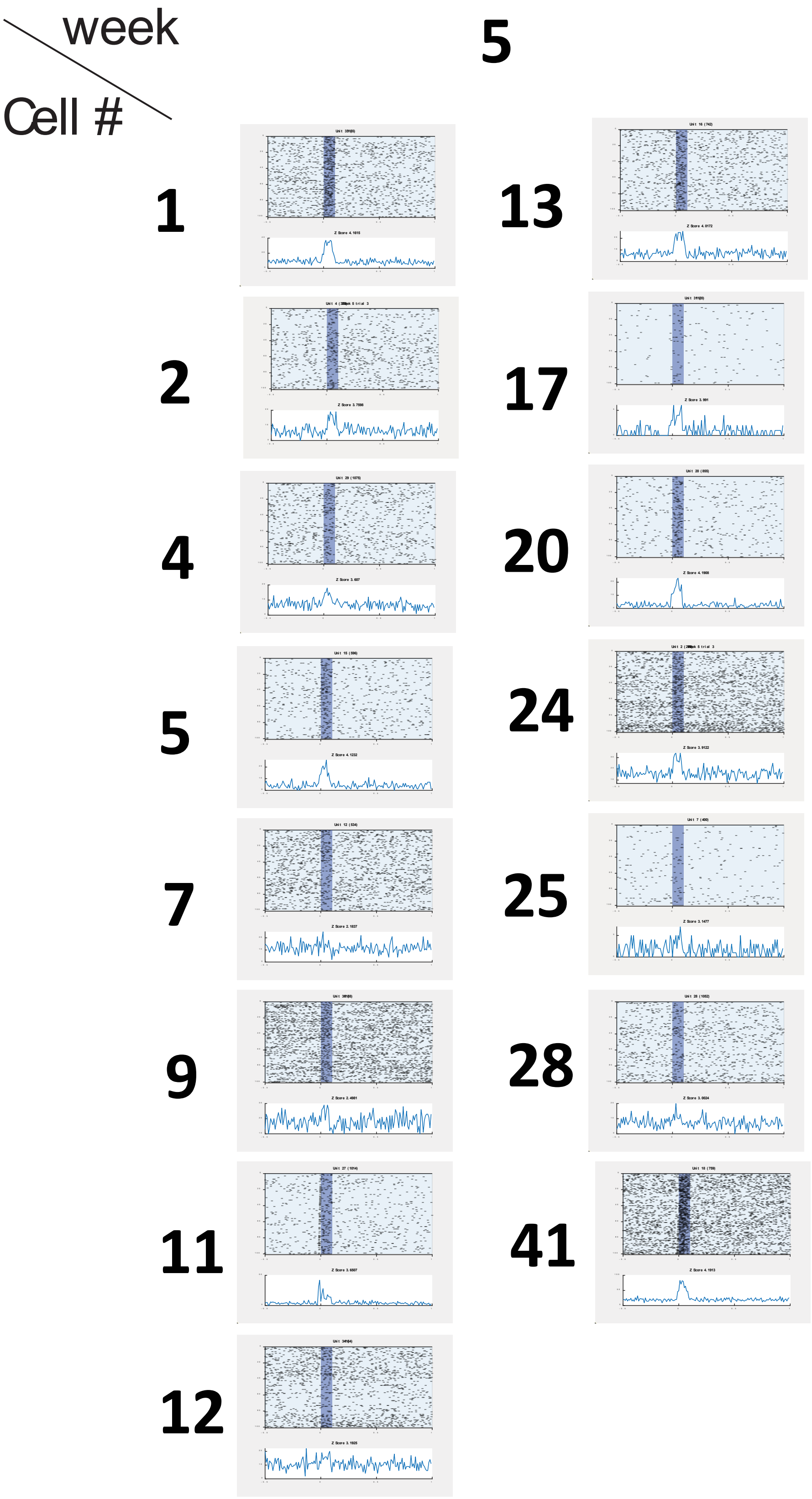
week
Cell #
5
1
2
4
5
7
9
11
12
13
17
20
24
25
28
41

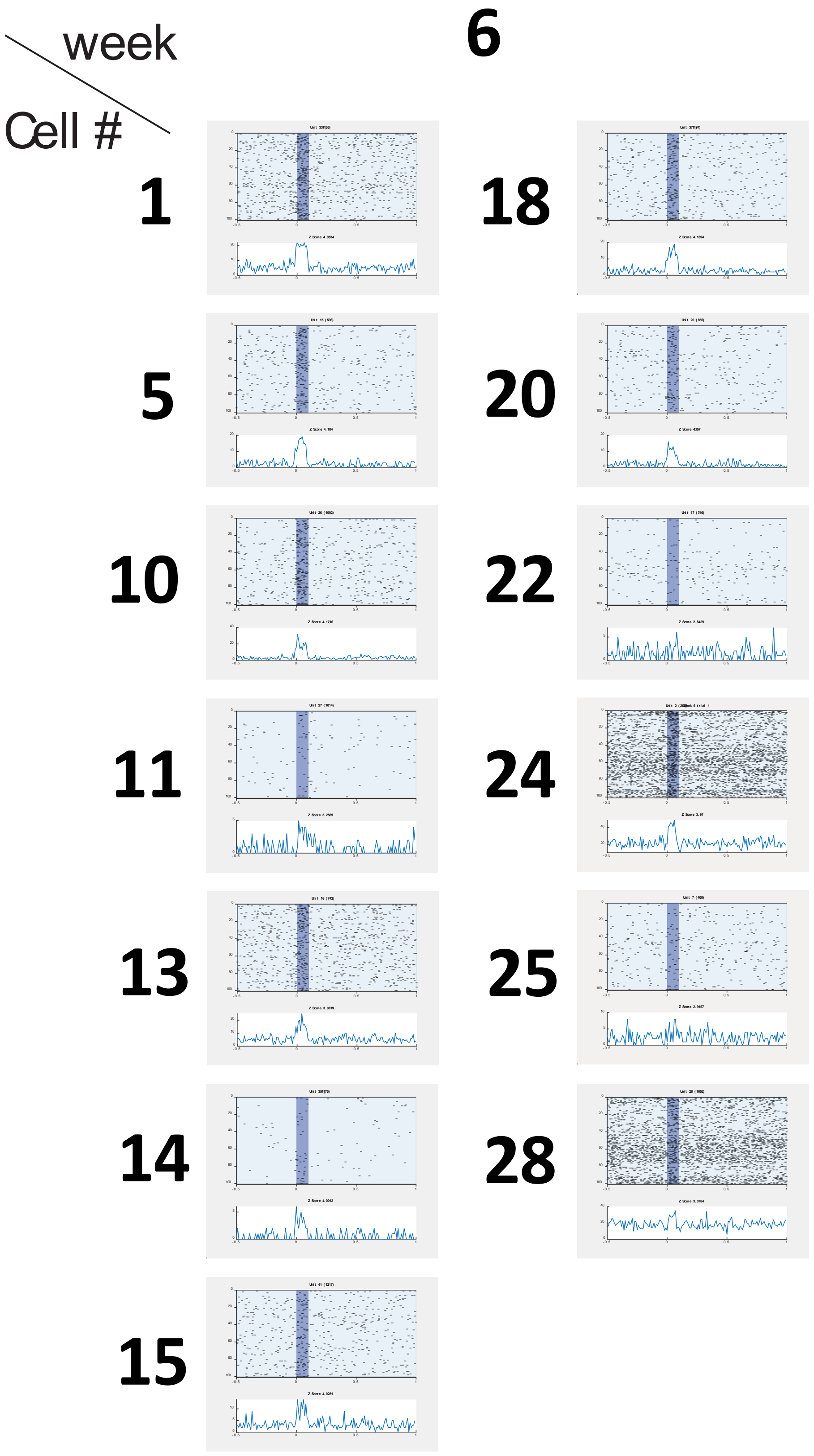

week
Cell #
6
1
18
5
20
10
22
11
24
13
25
14
28
15

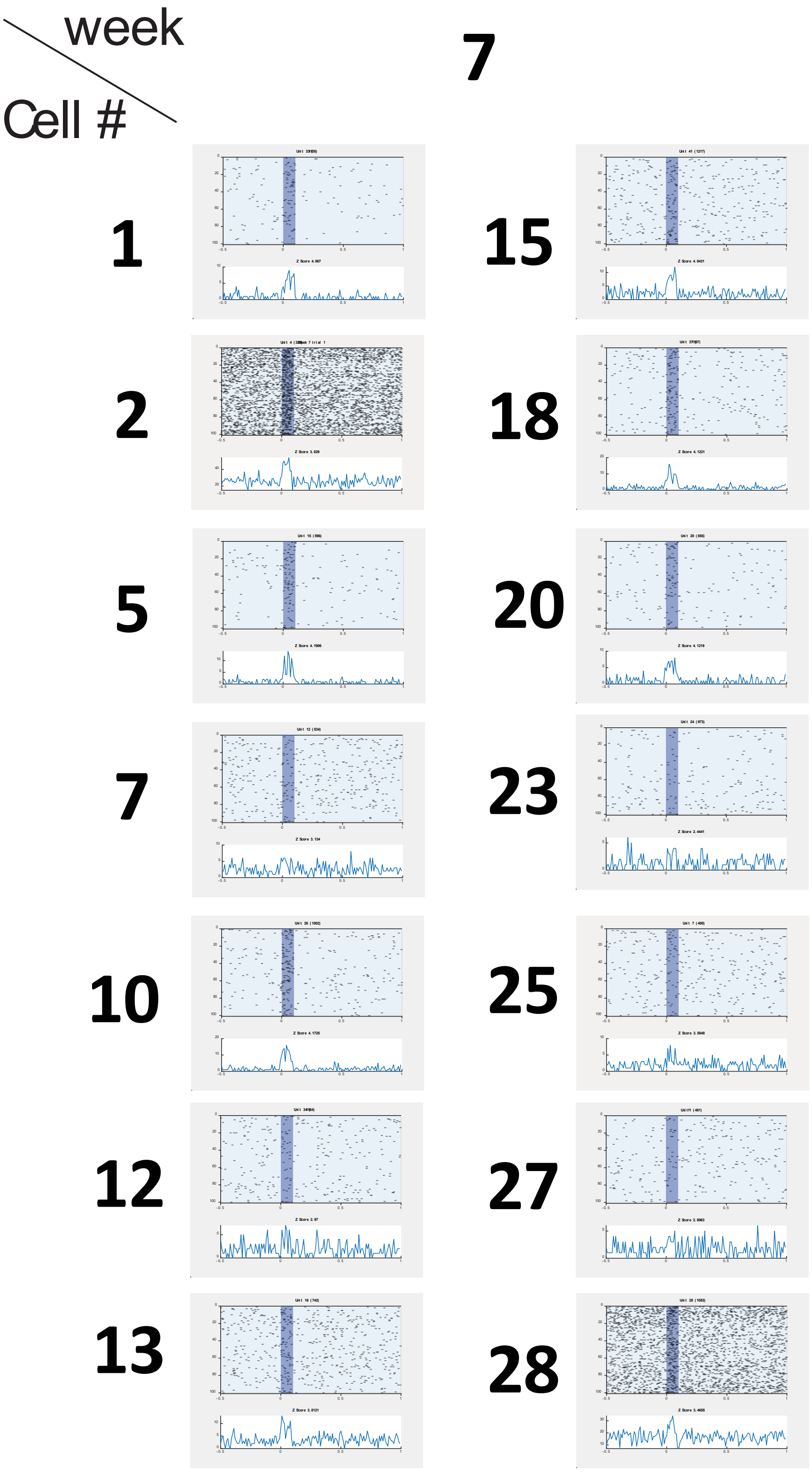
week
Cell #
7
1
15
2
18
5
20
7
23
10
25
12
27
13
28

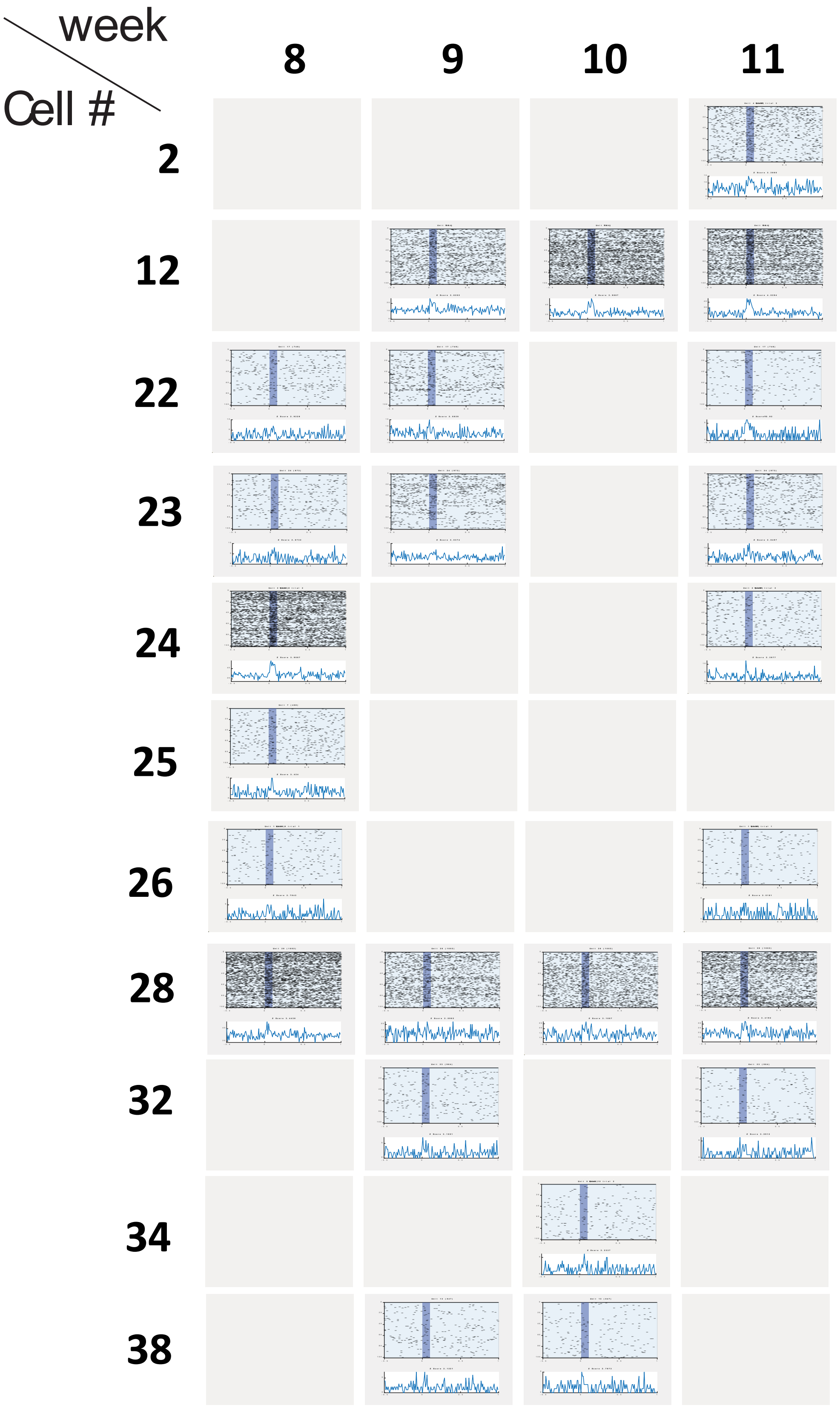
week
Cell #
8
9
10
11
2
12
22
23
24
25
26
28
32
34
38

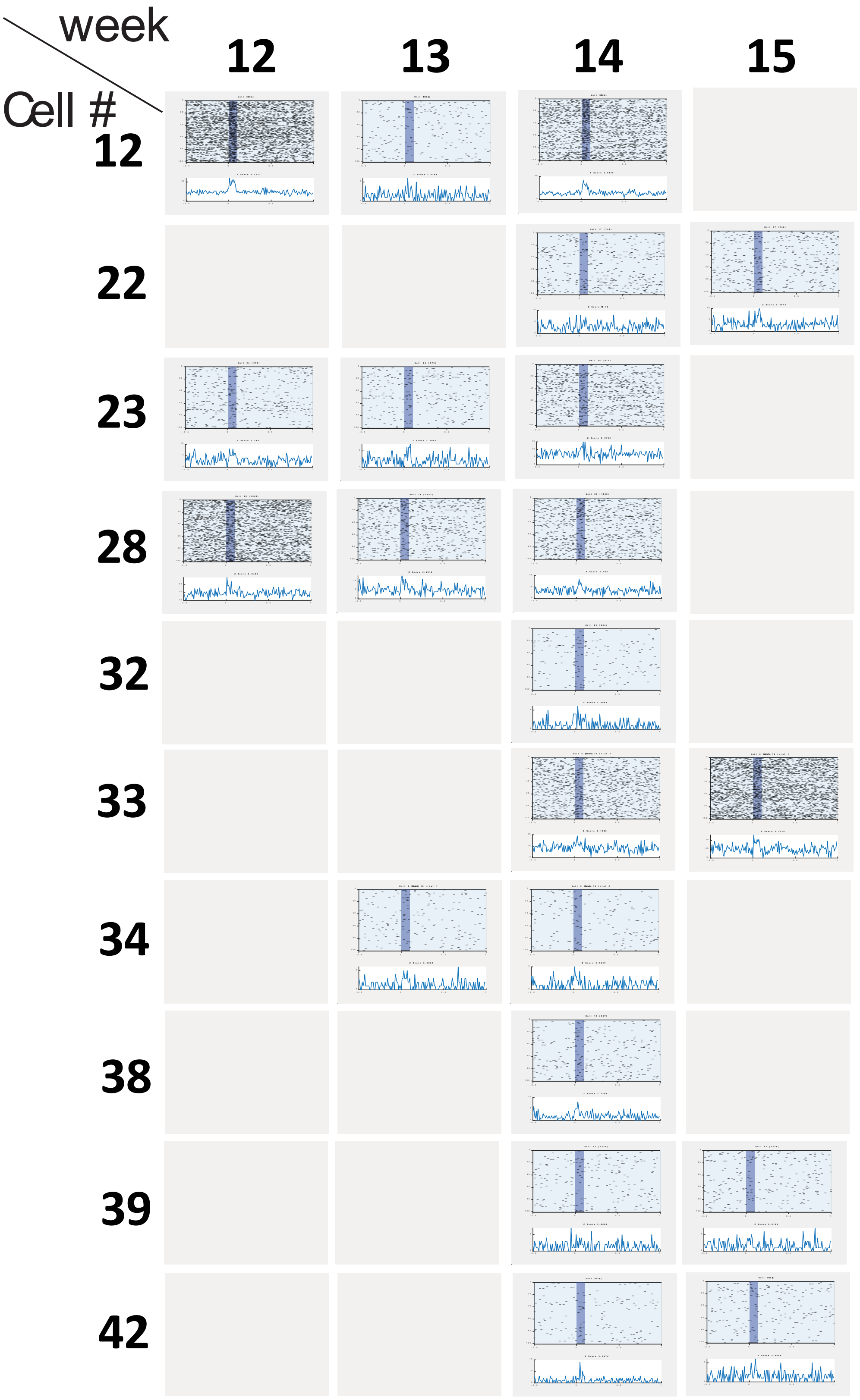
week
Cell #
12
13
14
15
12
22
23
28
32
33
34
38
39
42

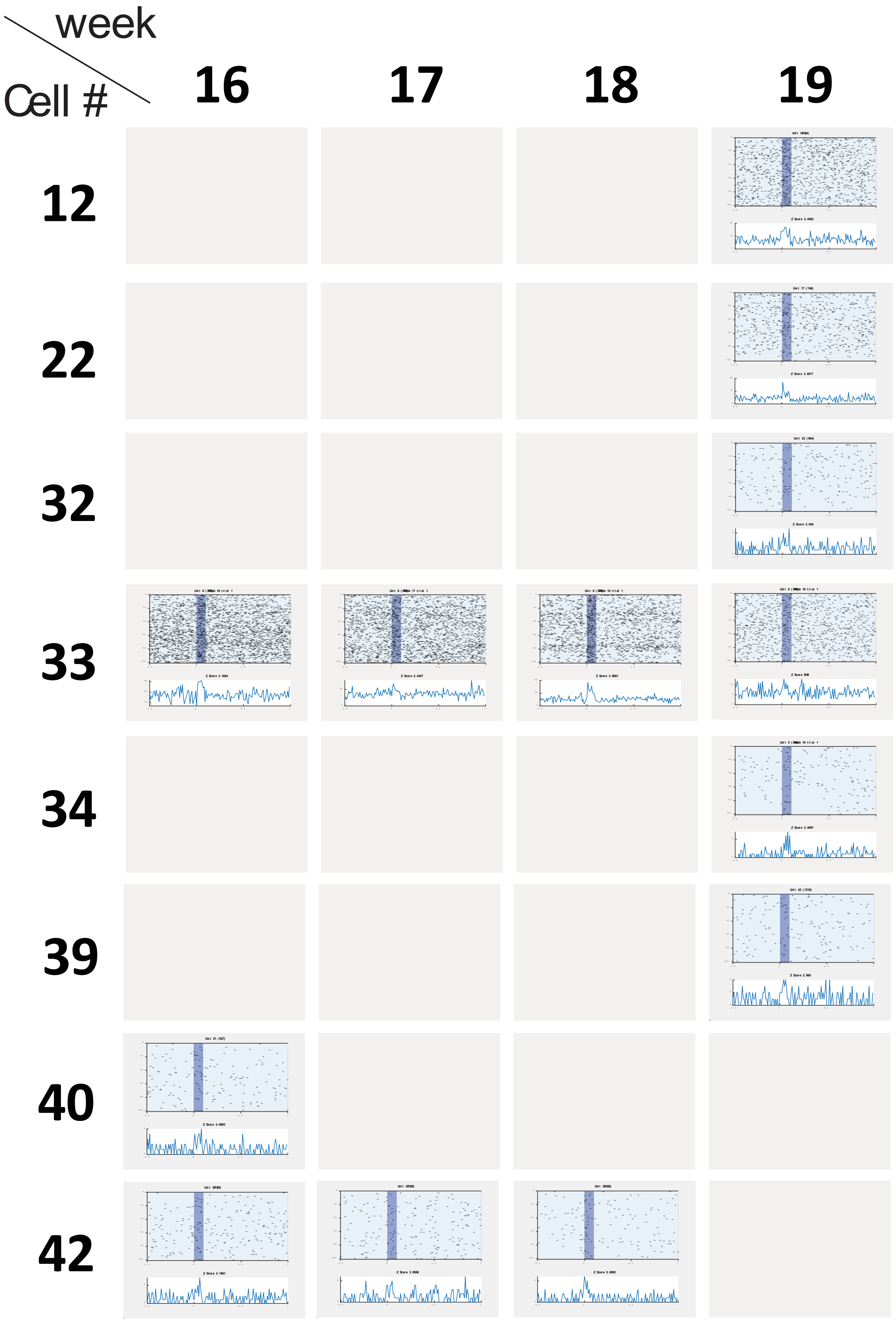
week
Cell #
16
17
18
19
12
22
32
33
34
39
40
42

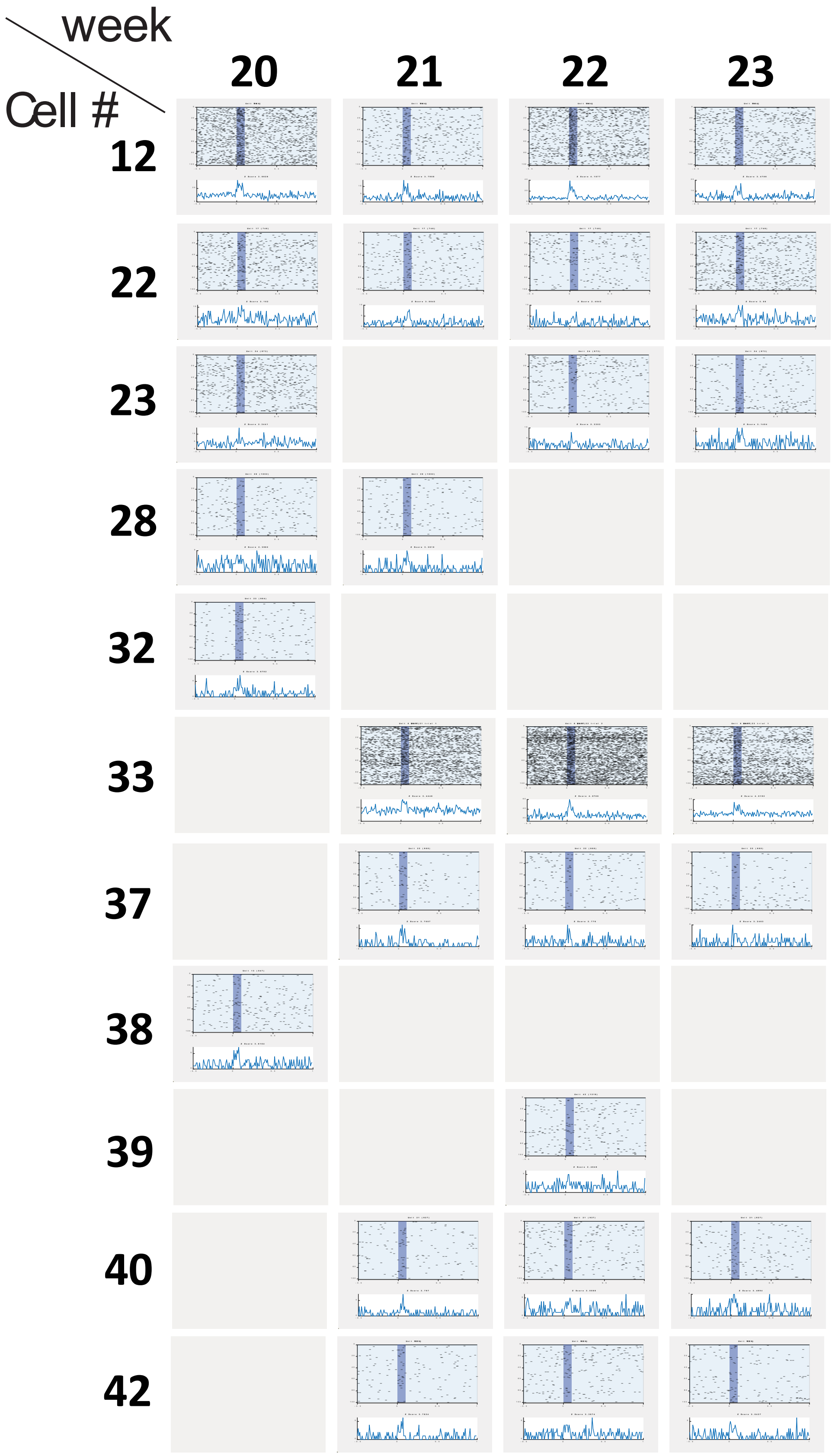
week
Cell #
20
21
22
23
12
22
23
28
32
33
37
38
39
40
42

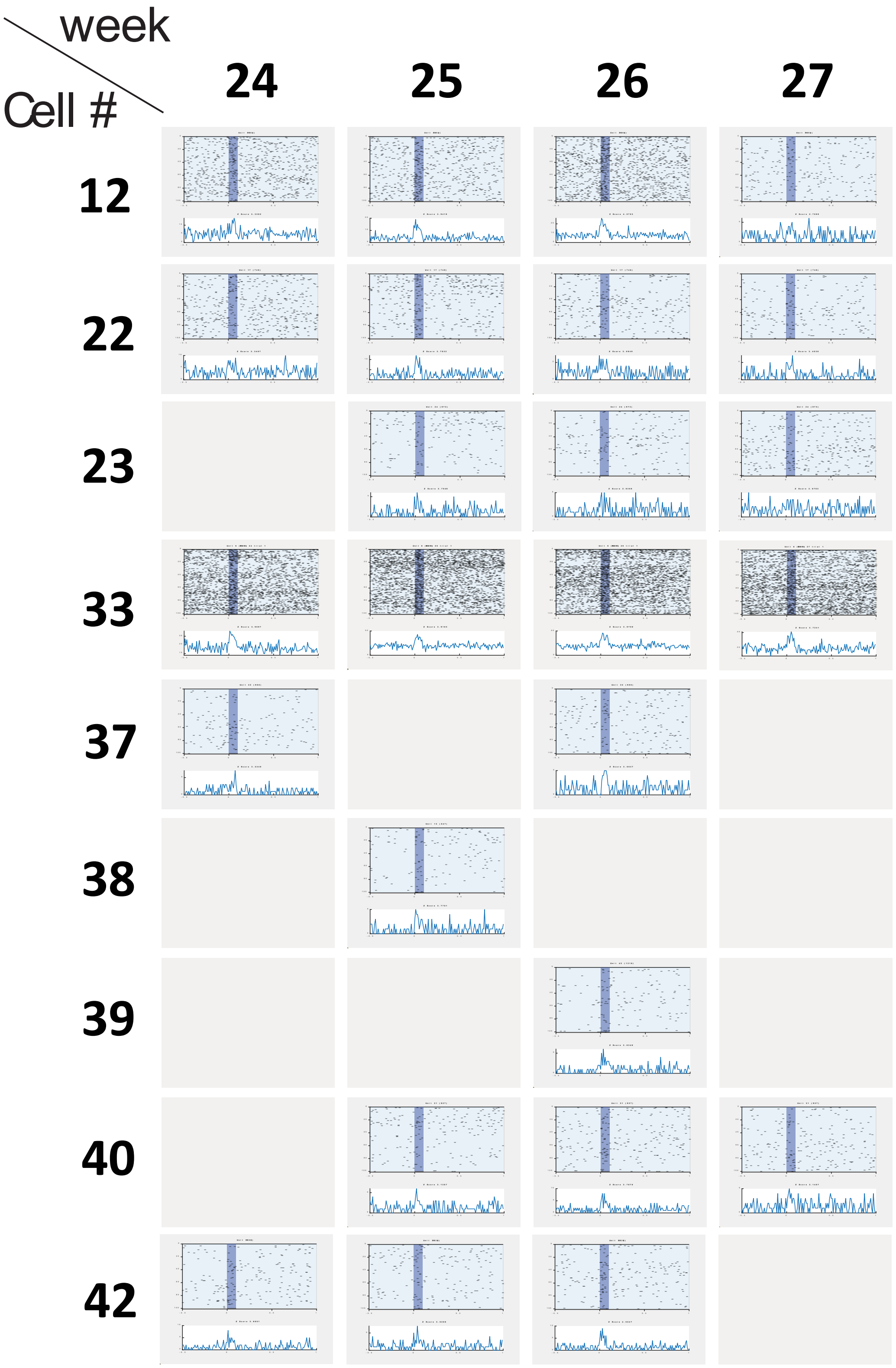
week
Cell #
24
25
26
27
12
22
23
33
37
38
39
40
42

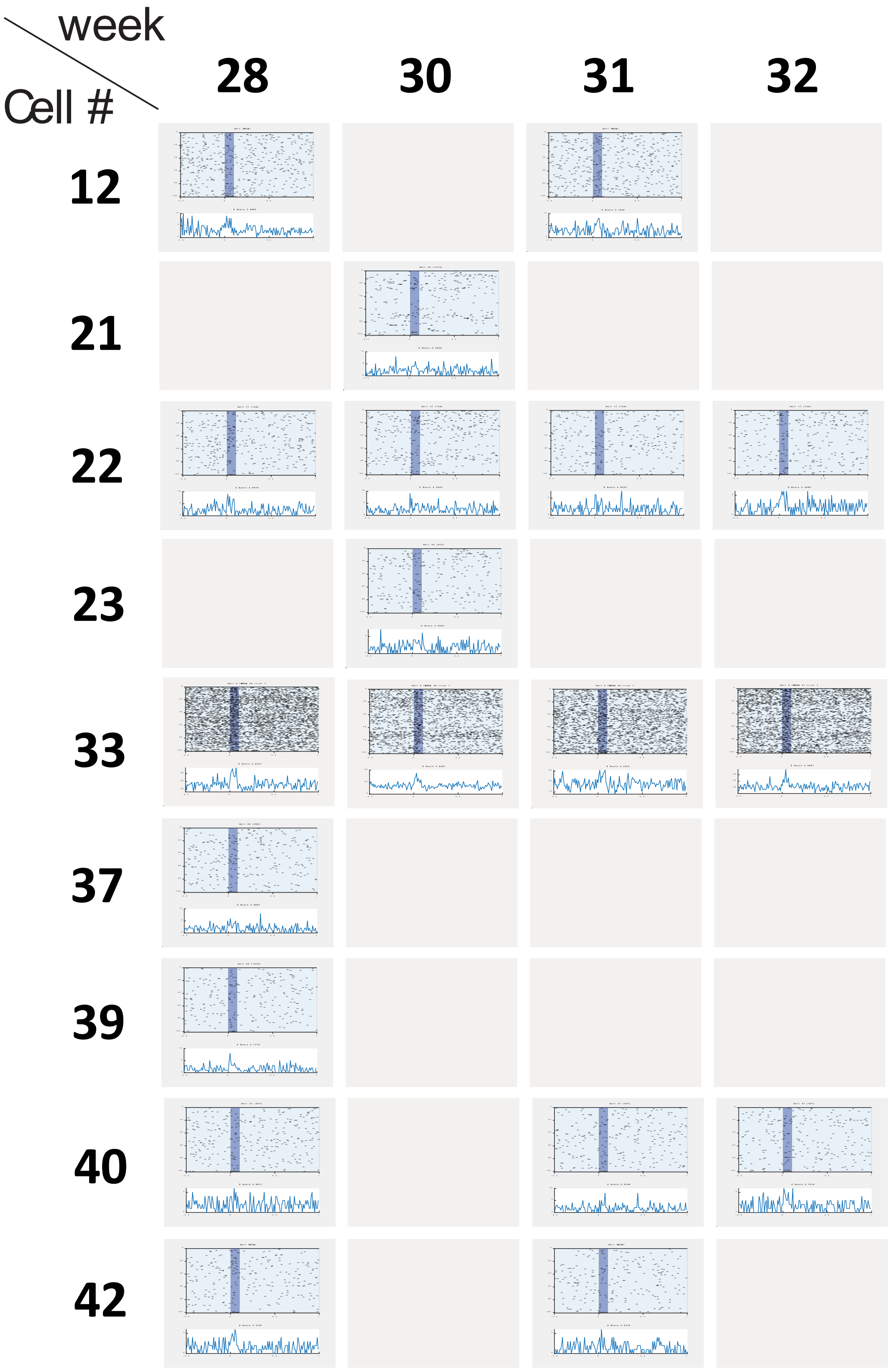
week
Cell #
28
30
31
32
12
21
22
23
33
37
39
40
42

**Supplementary Figure2.** Device 2 micro-LEDs I-V characterization after 13 months of implantation.

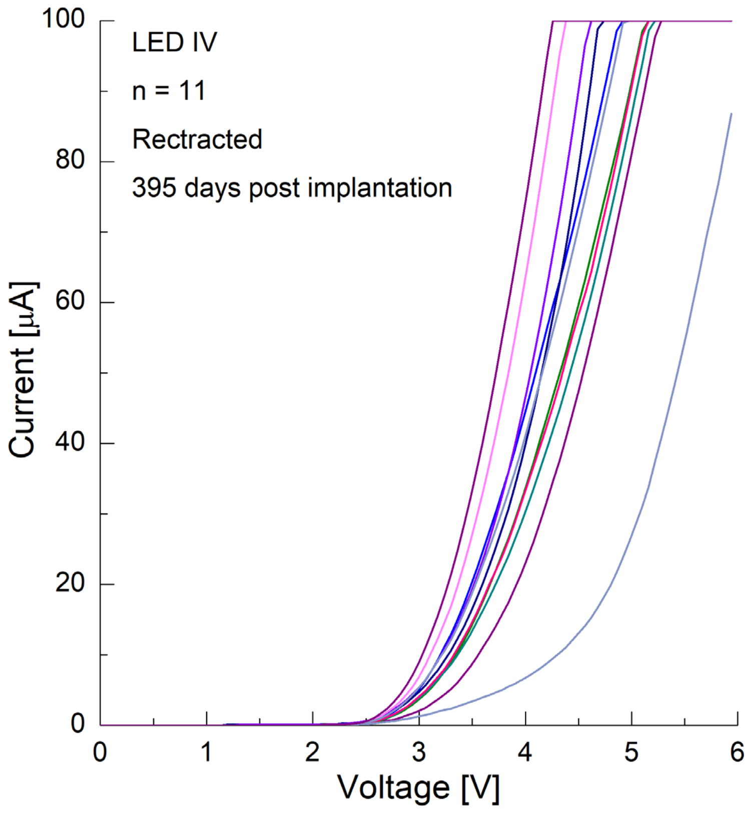

**Supplementary Figure3.** Explanted flexLiTE (top) and illumination of micro-LED after 13-month of implantation (bottom)

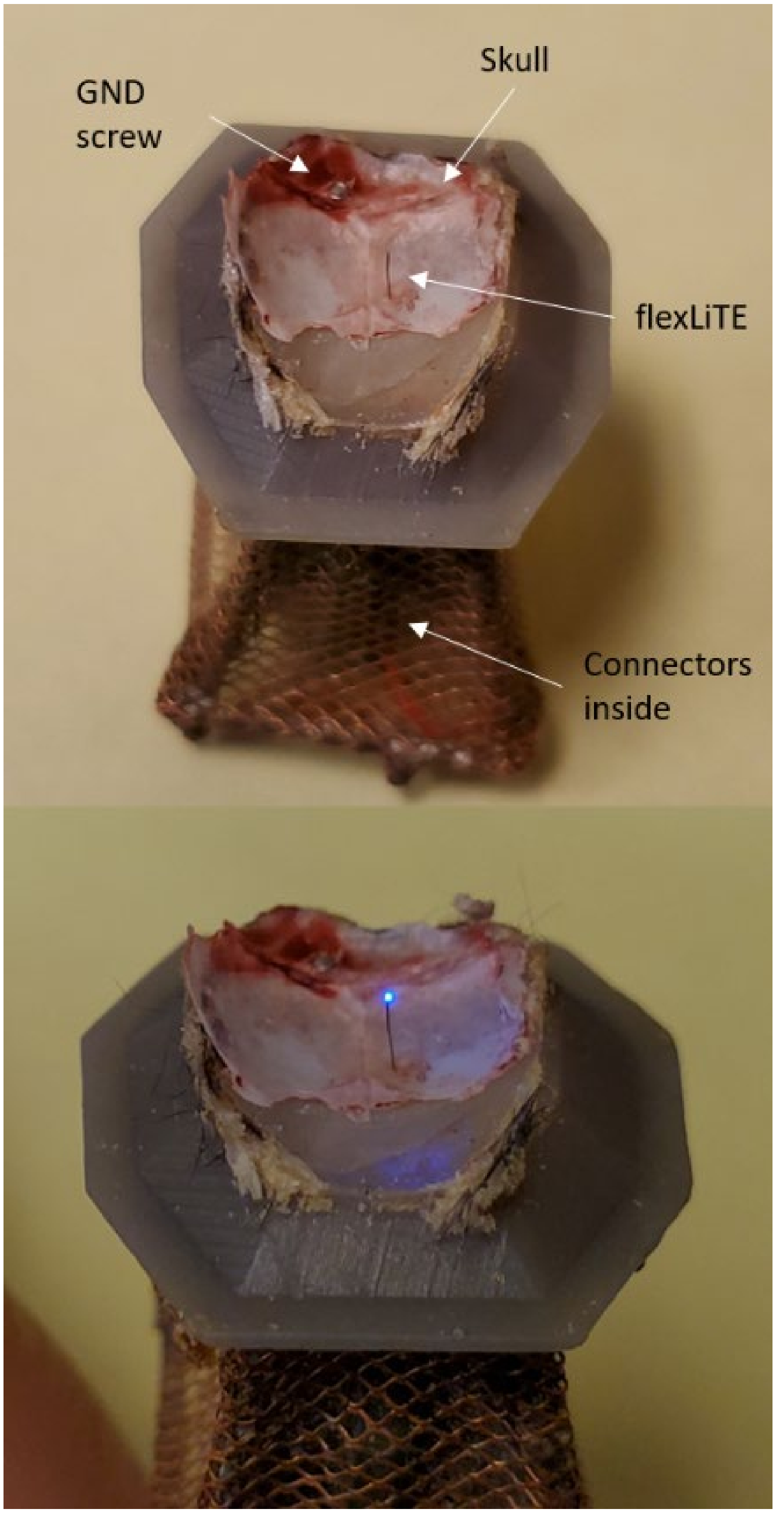
